# Engineering Quorum Quenching Acylases with Improved Kinetic and Biochemical Properties

**DOI:** 10.1101/2023.09.01.555929

**Authors:** Kitty Sompiyachoke, Mikael H. Elias

## Abstract

Many Gram-negative bacteria respond to *N-*acyl-L-homoserine lactone (AHL) signals to coordinate phenotypes such as biofilm formation and virulence factor production. Quorum-quenching enzymes, such as acylases, chemically degrade AHL signals, prevent signal reception by bacteria, and inhibit undesirable traits related to biofilm. These capabilities make these enzymes appealing candidates for controlling microbes. Yet, enzyme candidates with high activity levels, high substrate specificity for specific interference, and that are capable of being formulated into materials are needed. In this work, we undertook engineering efforts against two AHL acylases, PvdQ and MacQ, to obtain improved acylase variants. The engineering of acylase is complicated by low-throughput enzymatic assays. To alleviate this challenge, we report a time-course kinetic assay for AHL acylase that tracks the real-time production of homoserine lactone. Using the protein one-stop shop server (PROSS), we identified variants of PvdQ that were significantly stabilized, with melting point increases of up to 13.2 °C, which translated into high resistance against organic solvents and increased compatibility with material coatings. We also generated mutants of MacQ with considerably improved kinetic properties, with >10-fold increases against *N*-butyryl-L-homoserine lactone and *N-*hexanoyl-L-homoserine lactone. In fact, the variants presented here exhibit unique combinations of stability and activity levels. Accordingly, these changes resulted in increased quenching abilities using a biosensor model and greater inhibition of virulence factor production of *Pseudomonas aeruginosa* PA14. While the crystal structure of one of the MacQ variants, M1, did not reveal obvious structural determinants explaining the observed changes in kinetics, it allowed for the capture of an acyl-enzyme intermediate that confirms a previously hypothesized catalytic mechanism of AHL acylases.

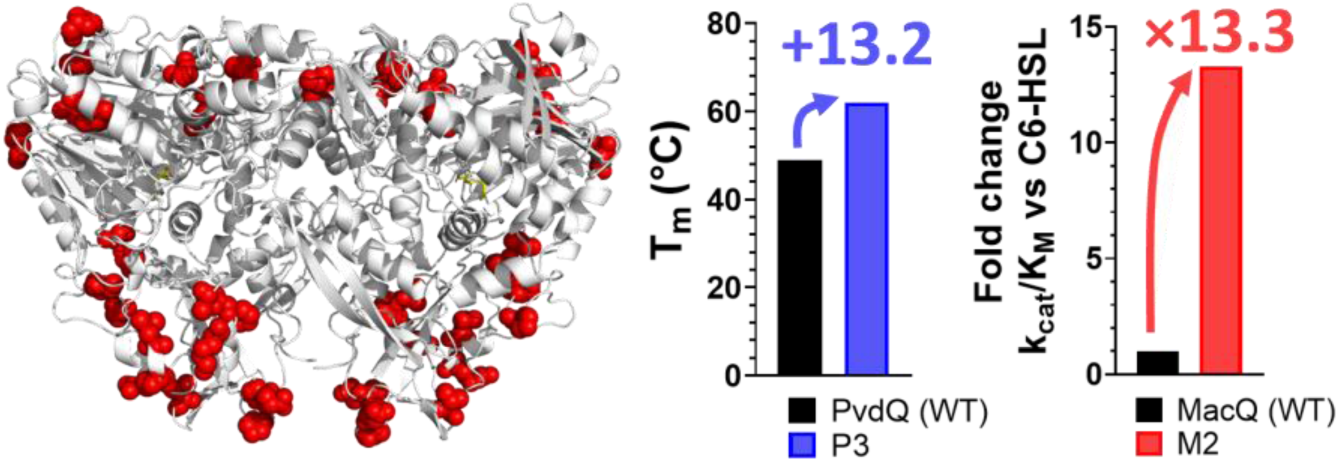

---

Bacterial autoinduction systems that are regulated by bacterial community cell density are known as quorum sensing (QS)^1^. QS systems are bacterial communication mechanisms, where constitutively produced small molecules accumulate with the increasing cell density and modulate gene expression profiles after reaching a threshold concentration. While the specific signaling molecule varies between bacterial species, many Gram-negative bacteria utilize various *N-*acyl-L*-*homoserine lactones (AHLs) as their autoinducer (autoinducer-I)^2^. AHL-based QS is used by numerous, clinically relevant, antibiotic-resistant pathogens such as *Pseudomonas aeruginosa*^3^*, Acinetobacter baumannii*^4^, all members of the *Burkholderia cepacian* complex^5,6^, as well as various environmental microbes such as *Chromobacterium violaceum*^7^, *Agrobacterium tumefasciens*^8^, or wastewater microbes^9^. The AHL signal binds to the cognate receptor (LuxR homologs), a cytoplasmic transcription factor that regulates a range of gene expression profiles^10^. This regulation and modulation of gene expression controls a myriad of bacterial behaviors, such as virulence, biofilm formation, and secondary metabolite synthesis^11,12^. The system is diverse and complex with redundancies and master regulators, where species can have more than one quorum sensing system that can feed into each other through regulation^11^.

AHL-degrading enzymes were found in bacteria and fungi^13^, and enzymes with AHL-degrading abilities were also found in mammals^14,15^. AHL-degrading enzymes may serve as a self-regulation measure for cells that use AHL signals to modulate the levels of self-cues, or as a defense measure to degrade other species’ cues in a shared environment^16^. Interestingly, AHL-degrading enzymes were previously shown to reduce bacterial QS-regulated phenotypes like biofilm formation and virulence in monocultures^17–21^ and complex communities^22,23^. The two main groups of AHL-degrading enzymes are lactonases (EC 3.1.1.81), which catalyze the hydrolysis of the lactone ring^17^, and acylases (EC 3.5.1.97), which cleave the amide bond of the AHL^24^.

Lactonases were shown to reduce virulence, biofilm formation and inhibit complex biological processes such as biocorrosion and biofouling^19,21,25–29^. Studies involving complex communities revealed that signal disruption by lactonases led to changes in the microbiome composition of an environment^22,28–30^. Similar experiments were conducted with AHL acylases and highlight similar ability to reduce biofilm and virulence, for example against *P. aeruginosa*, *Burkholderia cenocepacia*, and in wastewater communities^31,32^. Some AHL acylases, such as acylase I from *Aspergillus milleus*, PvdQ from *P. aeruginosa*, and porcine kidney acylase were formulated into materials such as polyurethane coatings, nanofibers, nanoparticles, and silicone as antibiofilm measures^33–40^. Remarkably, some microorganisms encode both AHL lactonases and acylases^41,42^. A major difference between the two classes of enzymes is the consequence of their different mechanisms: lactonases break open the lactone ring, while acylases cleave the amide bond. The action of acylases on AHLs results in the production of homoserine lactone (HSL) and the corresponding fatty acids. This difference might be important at low pH, where the AHL lactone ring opening was previously shown to be reversible^43^.

AHL acylases are N-terminal hydrolases that belong to the αββα-fold family and are homologous to penicillin acylases: while the fold itself is conserved, sequence conservation is low among known members of the family^44–46^. Characterized AHL acylases typically possess a broad substrate specificity spectrum, with preference for longer chain AHLs^47^. A major representative is PvdQ from *P. aeruginosa* PAO1, a quorum quenching enzyme that may also have roles in iron scavenging^48–50^. PvdQ prefers AHLs with an acyl chain greater than eight carbons in length^44,51^. A second AHL acylase was characterized from *Delftia sp*. VM4 and shown to be efficient at degrading medium length AHLs^52^. A third AHL acylase is MacQ from *Acidovorax* sp. MR-7 that was reported to possess a wide substrate specificity as shown by biosensors^53,54^, however its kinetic parameters against AHLs have not been reported to the best of our knowledge.

The AHL substrate preference of AHL-degrading enzymes may be important because the chemical structure of AHLs may confer specificity to bacterial signaling^55^. Previous engineering efforts to alter substrate specificity of quorum quenching enzymes have been performed, particularly on lactonases, facilitated by the availability and ease of activity assays^56–59^. Regarding AHL acylases, engineering efforts might be more challenging due to lower throughput enzymatic assays. Indeed, the AHL acylase activity has typically been measured using labor intensive methods such as gas chromatography^60^, HPLC analysis^61,62^, end-point assays^48,63^, and biosensor proxies^64–68^. Yet, previous efforts exist, including the alteration of the substrate specificity of PvdQ through rational design, which has resulted in a 3.8-fold decrease in affinity for a long chain substrate, 3-oxo-dodecanoyl-HSL, and a 4.3-fold increase in substrate specificity for *N-*octanoyl-HSL^34^.

In this study, we undertook engineering efforts for two AHL acylases, PvdQ and MacQ. We used the Protein Repair One Stop Shop (PROSS) tool that combines multiple sequence alignment, consensus scanning, and thermodynamic calculations to predict amino acid substitutions that favor stabilization of the overall protein structure^69^. Three different mutated sequences for each enzyme were produced and fully characterized. To determine the kinetic parameters of these enzymes against a variety of AHL substrates, we adapted a derivatization method for detecting the released homoserine lactone^64,70,71^ into a time course assay that can be reliably used for the kinetic parameter determination of AHL acylases. Stability of the acylase variants was examined and, while MacQ variants showed a reduction in melting temperature value (Tm), PvdQ variants showed increased thermal stability (T_m_>10 °C), consistent with the PROSS algorithm capabilities^69^. The increased stability of the PvdQ variants resulted in higher resistance in solvents and coatings, evidencing the importance of stability in efforts to formulate these enzymes into materials. Unexpectedly, some of the MacQ variants also showed altered substrate specificity profile with a 10-fold increase in catalytic efficiency for C4- and C6-HSL compared to the wild-type. The resolution of the crystal structure of variant M1 did not allow for determination of the specific determinants for this change but highlighted dynamical features that differ with the WT-MacQ. The structure captures an acyl-enzyme intermediate that complements previous insights on MacQ and confirms the catalytic mechanism for AHL hydrolysis.

## RESULTS AND DISCUSSION

### PROSS-generated MacQ and PvdQ variants

The acylase variants were created using the Protein Repair One-Stop Shop (PROSS) algorithm, which uses alignment scanning and computational mutation scanning to generate higher expressing and thermostable proteins^69^. For each acylase, namely MacQ and PvdQ, we produced the top three PROSS output sequences for the β-subunit (**table S1**). The three MacQ variants had 23, 29, and 32 amino acid substitutions making up 4%, 5.04%, and 5.57% of the total protein mutated. PvdQ had fewer mutations, being a smaller protein, with the three variants having 20, 26, and 29 substitutions each, composing 3.66%, 4.76%, 5.31% of the total protein. The majority of the substituted amino acids selected by PROSS retain their general properties such as hydrophobicity (**table S2**). The six variants M1, M2, M3 (for MacQ) and P1, P2, P3 (for PvdQ) were cloned and expressed as proenzymes as previously observed^53,62^. SDS-PAGE analysis after purification shows the presence of the two bands corresponding to both the α- and β- subunits, confirming that the autoproteolysis is successful (**figure S1**). We note that variant P3 showed a lower proportion of the processed subunits suggesting that the mutations might have interfered with the autocatalysis process. Expression levels of the MacQ variants were similar to wild type MacQ (WT-MacQ), while PvdQ variants expressed about 2- times less than wild type PvdQ (WT-PvdQ) in our experimental conditions (data not shown).

### Some acylase variants show substantially increased thermal stability

Unexpectedly, the MacQ variants exhibit reduced thermal stability. Indeed, they show T_m_ values reduced by 3.9, 8.3, and 12.6 °C for variants M1, M2, and M3, respectively (**figure 1A**). This may be because wild-type MacQ exhibit relatively high stability (T_m_ = 64.5 °C) and might be more stable than many of the acylase homologs used in the phylogenetic analysis performed by PROSS. The relatively high stability of wild-type MacQ was confirmed by determining the acylase activity temperature depende1ncy profile (T_m_ = 68.1°C; **figure S2**). On the other hand, many of the substitutions in the PvdQ variants are from polar uncharged amino acids to hydrophobic or charged amino acids. These changes may increase stability of the protein structure by improving packing of the protein structure and solvent-protein interactions to minimize enthalpy^72^. The WT-PvdQ enzyme exhibits lower thermal stability than WT-MacQ, as shown with the fluorescent dye-based thermal shift assay (T_m_ = 48.8 °C; **figure 1B**) and with the activity versus temperature dependency determination (T_m_ = 48.9 °C; **figure S2**).The generated PvdQ variants, P1, P2, and P3, show drastic increases in melting temperature compared to WT-PvdQ: 9.2, 11.7, and 13.2 °C for variants P1, P2, and P3, respectively (**figure 1B**).

**Figure 1:**
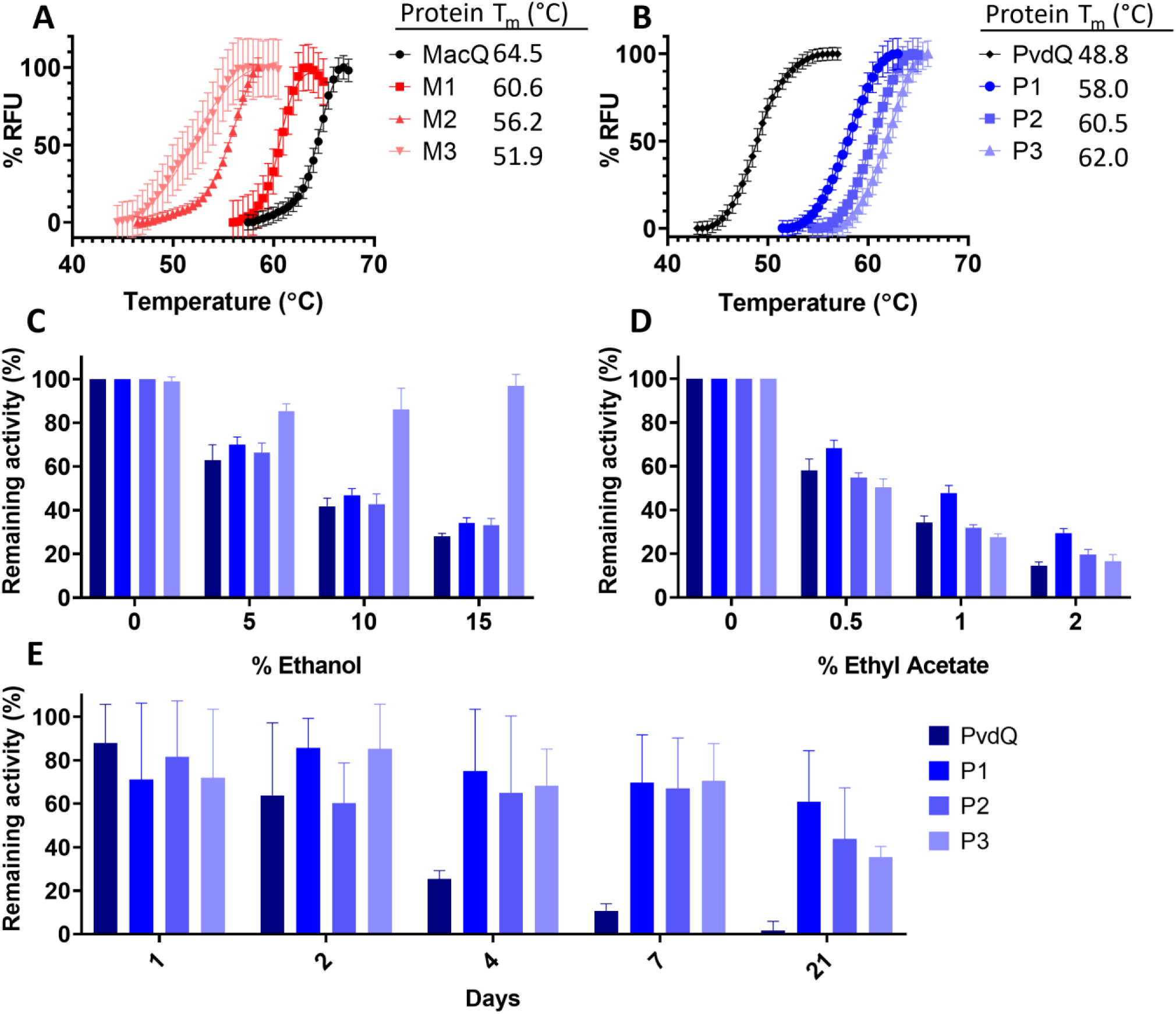
**Thermal and structural stability of acylases variants. A&B:** Melting curves ([**A**] PvdQ and its mutants; [**B**]: MacQ and its mutants) were determined using Sypro Orange as an indicator of protein unfolding over a range of tested temperatures. RFU: relative fluorescence units normalized to the maximum observed intensity. Measurements were performed in triplicates. **C&D**: Resistance of WT-PvdQ and variants P1, P2, and P3 in the presence of solvents. Activity levels were measured in C: ethanol and **D**: ethyl acetate against 3-oxo-C12-HSL. The final solvent percentage in the reaction mixture is shown as the x-axis. Measurements were performed in quadruplicates. **E**: Activity of PvdQ, P1, P2, and P3 acylases in dry silicon paint base over time. Shown is the remaining activity against 3-oxo-C12-HSL normalized to the activity in paint at day 1. Experiment was performed in quintuplicates.

### Thermostabilized PvdQ mutants have increased organic solvent resistance and are amenable to be formulated into material

We further characterized the stabilized PvdQ variants to determine how their observed increased melting temperature values relate to possible improvements in the enzyme’s compatibility with solvent or coating material. These tests were made possible using the adapted OPA assay and the substrate 3-oxo-C12-HSL (see methods). The stability of the PvdQ mutants was probed by measuring their activity in the presence of increasing concentrations of ethanol and ethyl acetate (**figure 1C, D**). These solvents can be used in adhesives, paints, and coatings: resistance to these solvents is a desirable trait for an enzyme with potential to improve materials. Consistent with their increased T_m_ values, all mutants outperformed WT-PvdQ in all tested ethanol concentrations. The P3 mutant also retains over 80% original activity in the presence of 15% ethanol and showed >2-fold higher activity than WT-PvdQ and the other variants in this condition (**figure 1C**). In ethyl acetate, however, differences between WT-PvdQ and the variants were less marked. Yet, P1 showed the highest resistance to ethyl acetate (**figure 1D**).

We continued our evaluation of the stability of the variants by probing their activity in a silicone coating base (**figure 1E**). Results show that the PvdQ mutants drastically outperformed their wild-type counterpart. All variants maintained >60% of their original activity after 4 days of storage as dried silicon paint, while WT-PvdQ lost ∼70% of its original activity. After 21 days, variant P1 retained ∼60% of activity, while the WT-PvdQ containing coating was nearly inactive. These results suggest that the thermostabilized variants may have improved compatibility with solvents and coatings than the wild-type enzyme. Therefore, they might represent appealing candidates for the creation of new materials with antibiofilm activity, or for the study of interference in QS in conditions unfavorable to less stable enzymes.

### A time-course kinetic assay for quorum quenching acylases

Acylase activity has rarely been monitored in real-time, but rather through HPLC analysis, observing and quantifying the release of the fatty acid and free homoserine lactone through OPA-derivatization^13,61,71^ and/or mass spectrometry^61^. Several end point kinetic assays were previously reported, including using o-phthalaldehyde (OPA) as an end-point indicator of acylase activity, where it is added after reaction termination and the absorbance values compared to a calibration curve with or without chromatographic separation^34,64,73^. A similar method is using fluorescamine as the fluorogenic indicator of homoserine generation^74^. However, these approaches are time consuming and low throughput when measuring kinetic parameters for multiple substrates and make acylase engineering more challenging.

Here, we adapted the OPA assay and established a time course protocol. The homoserine lactone produced by the AHL acylase reaction reacts with OPA and dithiothreitol to form a fluorescent isoindole product (**figure S3**)^75^. We established conditions where the increase in product formation as shown by fluorescence is enzyme concentration-dependent (**figure S4**). While OPA may react with free amines at the surface of the proteins, we noticed that the (i) background is low and (ii) the possible inhibition of the tested enzymes is minimal during the time course of the reaction. Background reaction was taken into account by an initial incubation of the enzyme with OPA for 5 minutes, prior to starting the enzymatic reaction by adding substrate AHL.

We validated this time-course assay by first determining the kinetic parameters for the WT-PvdQ (**table 12**) and found that the determined values are very close to the previously reported values using the OPA end-point assay^34,51^ (**table S3**). For example, previous studies reported catalytic efficiencies for WT-PvdQ that were relatively consistent, within a ∼2-fold range: 2.3 x 10^3^ s^-1^ M^-1^ and 5.8 x 10^3^ s^-1^ M^-1^ against 3-oxo-C12-HSL ^34,51^, and 2.2 x 10^2^ s^-1^ M^-1^ and 0.8 x 10^3^ s^-1^ M^-1^ for C8-HSL^34,51^. The catalytic efficiency values determined in the present study are in good agreement with these, with 2.22 x 10^4^ s^-1^ M^-1^ against 3-oxo-C12-HSL and 1.75 x 10^3^ s^-1^ M^-1^ against C8-HSL, or ∼2-fold higher than the values reported by Koch et al. (2014)^34^. This confirms the validity of the values determined with this method that allows monitoring of the time-course reaction of AHL degradation by acylase using fluorescence.

**Table 1:**
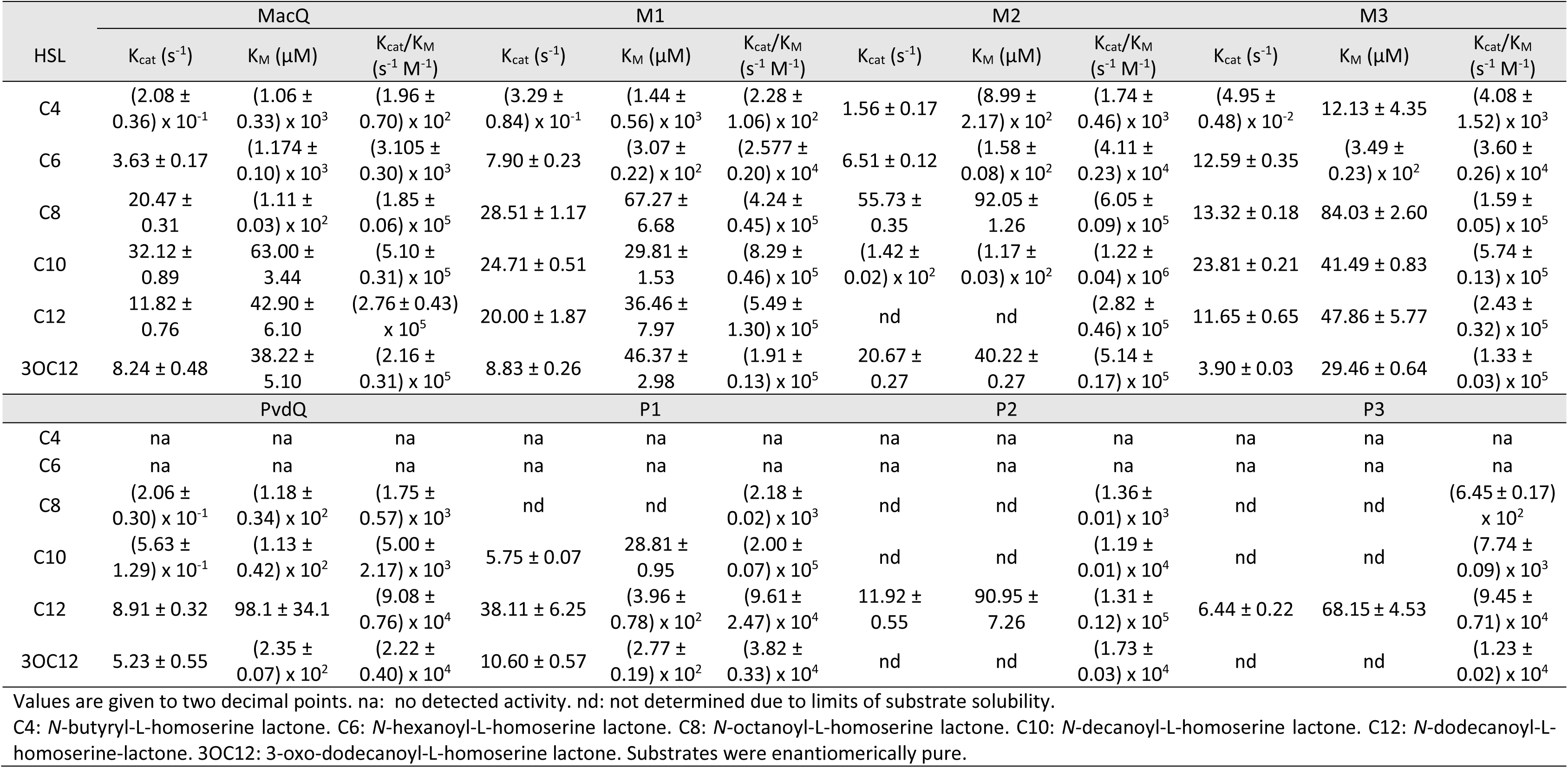
Kinetic parameters against N-acyl-L-homoserine lactone substrates for MacQ and PvdQ acylases and their mutants.

### Some PROSS-generated mutants show altered substrate specificity

WT-PvdQ shows activity against AHLs with acyl chains between eight to twelve carbons long. The variants exhibit similar substrate preference profile and overall catalytic efficiency levels to WT-PvdQ. Yet, it appears that their K_M_ values have increased. This is evidenced by the observation that saturation could not be reached for some substrates (**table 1, figure S5-S6**). Of note, the three variants show higher catalytic efficiencies than WT-PvdQ against C10-HSL, by 40-fold, 2.4-fold, and 1.5-fold, for P1, P2, and P3, respectively.

The change in substrate specificity was more drastic for MacQ mutants. The mutants were generally more active than WT-MacQ on short chain AHLs (**table 1**). This is true for C8-HSL for which catalytic efficiencies of M1 and M2 are 2.3-fold and 3.3-fold higher than WT-MacQ, respectively. These changes are stronger for C6-HSL where M1 shows catalytic efficiencies 8- fold higher than WT-MacQ and the other variants have >10-fold increases than WT-MacQ. Similarly, M2 exhibits ∼9-fold higher catalytic efficiency and M3 has ∼21-fold higher catalytic efficiency against C4 HSL as compared to the WT enzyme (**table 1**). These changes in substrate specificity are significant in the context of AHL acylase engineering. To illustrate this, previous engineering efforts on PvdQ resulted in ∼4-fold changes in catalytic efficiencies against C8- and C12-HSLs^34^. These results are therefore evidence for the potential of PROSS to generate functionally diverse mutants.

The properties of the generated variants of MacQ were further examined using a biosensor assay. This biosensor consists of an *E. coli* strain expressing GFP under control of the LuxI/R system that responds to C6-HSLs (**figure 2A**)^65,68^. With this assay, both M1 and M2 were more effective at completely quenching the C6-HSL signal than WT-MacQ, with complete quenching observed after ∼70, ∼80 and ∼120 min for M2, M1, and WT-MacQ, respectively (**figure 2B**). Unexpectedly, despite higher catalytic efficiencies, M3 only performed as well as WT-MacQ, possibly due to its reduced stability as measured by its lower melting temperature (**figure 1**).

**Figure 2:**
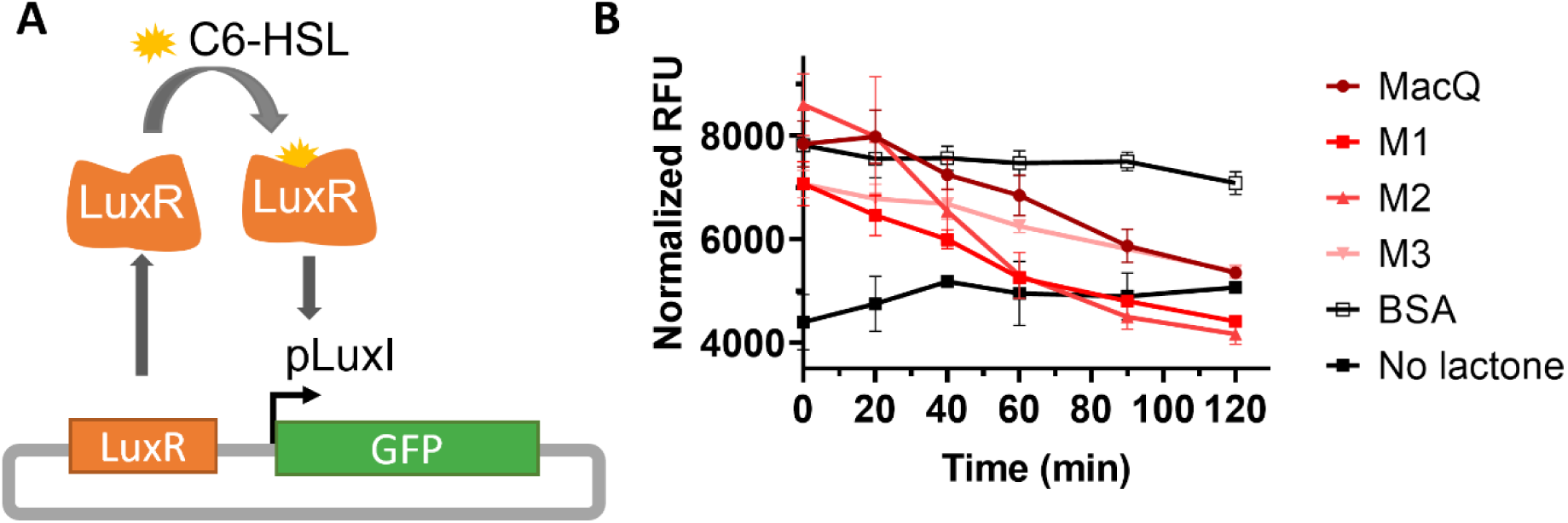
**A:** Schematic of LuxI/R based biosensor plasmid pJBA132 controlling the expression of green fluorescent protein (GFP) upon the binding of *N*-hexanoyl-L-homoserine lactone (C6-HSL) to pLuxI. **B:** GFP fluorescence of *E. coli* expressing pJBA132 after induction with C6-HSL incubated with acylases for the set amount of time. DMSO was used as the background control and bovine serum albumin (BSA) used as a negative control. Experiments were performed in quadruplicates.

### Engineered acylase variants are effective at reducing biofilm formation and virulence factor production in PA14

We evaluated the ability of MacQ and PvdQ variants to quench *P. aeruginosa* PA14 (PA14) cultures. PA14 is known to utilize two parallel AHL-based QS circuits relying on C4- HSL and 3-oxo-C12 HSL^76^ that are involved in biofilm formation and virulence factor production. We thus grew PA14 cultures overnight in the presence of our acylase variants and performed virulence factor production assays to assess biofilm formation and phenotype of PA14 after treatment (**figure 3**).

**Figure 3:**
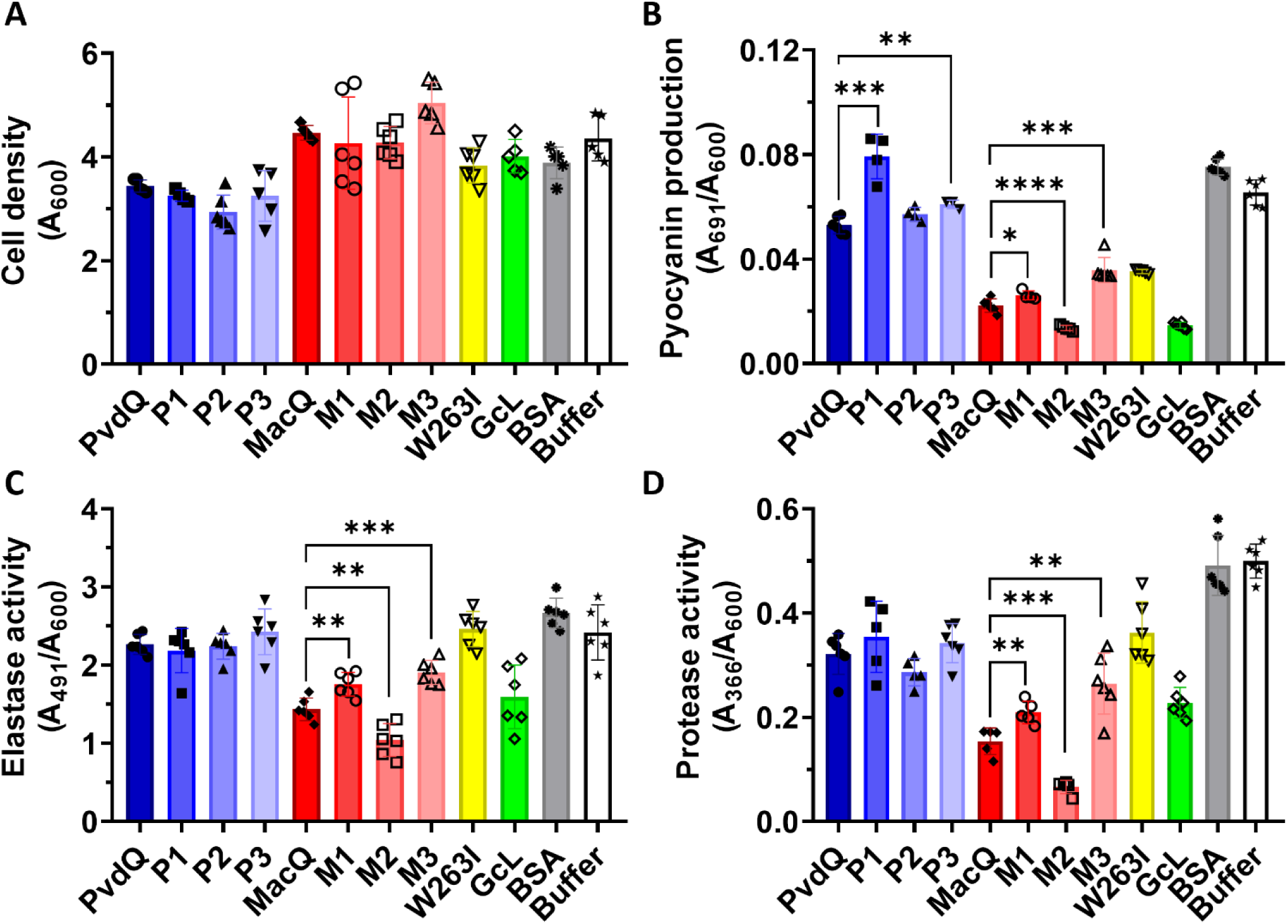
Growth and virulence factor production analysis of *P. aeruginosa* PA14 after treatment with acylases and lactonases. PvdQ, P1, P2, P3, MacQ, M1, M2, M3: acylases tested in this study. Lactonases SsoPox W263I and GcL are used as positive controls and benchmark. Bovine serum albumin and Buffer (50 mM HEPES pH 7.0, 150 mM NaCl, 0.2 mM CoCl_2_) are used as negative controls. **A**: Cell density measured at A_600_. **B**: pyocyanin in the cell supernatant measured at A_691_, **C**: elastase activity in the supernatant measured through Congo Red released from elastin-Congo Red measured at A_491_, **D**: protease activity in the supernatant measured through release of azo dye from azocasein, normalized to cell density. Statistical significance was determined using an unpaired t-test. * P .0.5; ** P .0.01; ***P .0.001; **** P .0.0001. N = 4-6 replicates.

Consistent with previous reports^48,53^, both WT-PvdQ and WT-MacQ are effective at reducing virulence factor production without inhibiting cell growth in bacterial pathogens that use AHLs such as PA14. This is also true for the variants generated in this study (**figure 3A**).

With regards to MacQ variants, M2 showed the highest inhibitory activities, with 78%, 56%, and 84% inhibition for pyocyanin, elastase and protease activity, respectively, compared to the buffer only control (**figure 3B-D)**. This increased inhibitory effect is consistent with the variant’s increase in catalytic efficiency against both PA14’s signaling molecules, with >8-fold improvement in catalytic efficiency against C4-HSL and >2-fold higher efficiency against 3- oxo-C12-HSL, as compared to the other variants and WT-MacQ. M1 only shows slightly higher catalytic efficiency against C4-HSL and lower activity against 3-oxo-C12-HSL. A similar pattern is seen for M3. For PvdQ variants, results show little variation in their ability to inhibit elastase and protease activity (**figure 3C, D**), and overall, the variants show WT-PvdQ-like quorum quenching activity. This is consistent with their overall similar kinetic parameters to WT-PvdQ (**table 1**).

This study is also an opportunity to compare the inhibitory effects of different quorum quenching enzymes on PA14, namely acylases and lactonases. To the best of our knowledge, this comparison has not been reported in prior reports. Here we compare WT-MacQ, WT-PvdQ and their variants with the lactonases GcL, a broad spectrum enzyme^77^, and SsoPox W263I, a mutant of SsoPox with preference for long AHLs^57^. Results show that MacQ and its variants were more effective at quenching the measured virulence factors (biofilm, pyocyanin and proteases). Its catalytic rates are 10-fold higher than those of PvdQ on long-chain AHLs and, in addition, MacQ is capable of degrading shorter chains, whereas PvdQ is not (**table 1**). A similar observation could be made with the two lactonases: the generalist GcL showed greater quorum quenching activity than SsoPox, which prefers long AHLs (**figure 3**). Additionally, MacQ and its mutants were as or more effective at reducing virulence factor production than the lactonases tested in this experiment, including the highly active GcL lactonase^78^. Because WT-MacQ and GcL show similar catalytic efficiencies, the observed differences might evidence complexity in the regulation and mechanisms of quorum quenching in the cell, such as inhibition occurring through different signaling cascades, that have yet to be elucidated.

### Variant M1 exhibits subtle structural changes

To investigate the origins of the differences in properties between the wild type acylases and the variants, we attempted to crystallize the different variants reported here. We could only produce diffracting quality crystals for MacQ variant M1 (**figure 4, table S4**). The structure of M1 highlights the location of the PROSS-generated mutations: at the periphery of the protein, on the surface and away from the active site (**figure 4B**). This likely allows the variants to retain their AHL-acylase activity.

**Figure 4:**
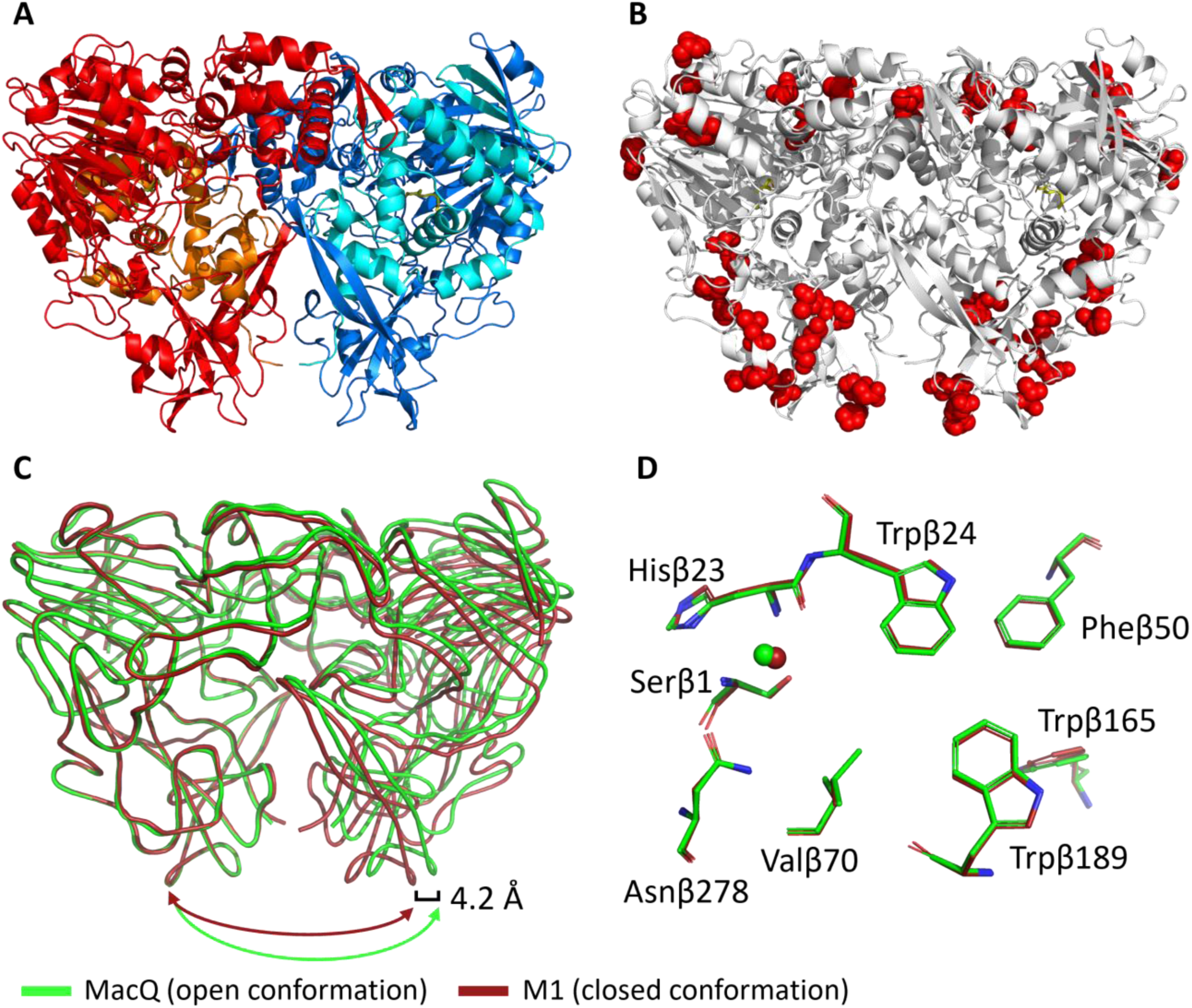
Crystal structure of M1 (PDB: 8S05). **A**: Homodimer of heterodimers. Red and blue: β subunits, orange and cyan: α subunits. Dodecanoic acid ligand is shown in yellow. **B**: Mutation sites in M1 are shown as red spheres. **C**: alignment of M1 (red) with MacQ (green, PDB: 4yfa) showing smoothed loop positions. **D**: active site alignment of M1 (red) and MacQ (green). Residue numbering refers to M1.

The structure of M1 is overall very similar to the WT-MacQ structure (**figure 4C**). However, we do note some differences in loop positions and the possible rotamer conformations. In particular, the dimer of heterodimers configuration shows a more “closed” configuration as compared to WT-MacQ. The structure closes by ∼4.2 Å and the second heterodimer appears translated by ∼3 Å (**figure 4C**). These changes in the enzyme configuration might result in different protein dynamics and contribute to the observed changes in catalytic efficiencies.

In addition to these global configurational changes, smaller, local different conformations are observed throughout the M1 structure (**figure S7**). Some of these differences appear directly related to the mutations (**figures S7A, B**), for example, the Serβ333Pro appears to disrupt an α-helix (**figure S7D**). Another example related to secondary structure movement are the mutations Metβ227Lys, Valβ229Phe, and Glyβ230Ala affect the α-helix and alter the backbone position by ∼2 Å (**figure S7B**).

These conformational differences may contribute to the observed reduction in melting temperatures (-3.9 °C) and increase in catalytic efficiency (>8-fold against C6-HSL) of M1 as compared to WT-MacQ. We note that the obtained M1 structure does not show electron density corresponding to some maturation peptide fragments (spacer peptide), contrary to the structure of WT-MacQ^54^. While fragments originating from AHL acylase maturation were reported to be unnecessary for activity^48,61^, this absence could also contribute to some of the observed conformational differences between the two structures. Another potential reason for the observed different conformation is that the structure of WT-MacQ shows only one covalent acyl intermediate, while the M1 structures shows it for both catalytic domains. Overall, the active site configuration of the M1 variant is very similar to that of WT-MacQ (**Figure 4D**), confirming that our engineering approach leaves the active site and chemistry around the site intact.

### The structure of variant M1 reveals a captured acyl-enzyme intermediate

The examination of the electronic density maps reveal density corresponding to a covalently bound adduct on the catalytic serine Serβ1 (**figure 5A**). This was modeled as decanoic acid based on the electronic density map (**figure 5A**). Because no substrate was added during crystallization, we hypothesize that it may originate from the *E. coli* host but can only speculate on the nature of the molecule that reacted with the enzyme. To illustrate this possibility, WT-MacQ was previously shown to be a promiscuous enzyme that for example exhibit β-lactamase activity^53^, and may therefore react with unknown cellular molecules. The capture of a similar acyl intermediate was previously reported for C10-HSL soaked WT-MacQ (PDB: 4yfa)^54^ and C12-HSL soaked WT-PvdQ (PDB:2wyb)^44^ (**figure S8B, C**). As expected, the structure of M1 aligns very closely to the structure of WT-MacQ (**figure S8D**). Minor conformational changes of Serβ1, Hisβ23 and the recycling water molecule can be observed. A main difference is the rotation of Hisβ23 sidechain, of about 0.4 Å towards the active site Serβ1. The active site of WT-PvdQ shows some differences in the residues lining the active site, resulting in a slightly different binding mode, particularly with regards to the acyl chain (**figure S8C, E**). For example, Trpβ24 adopts a slightly different conformation than the equivalent residue in WT-PvdQ (Pheβ24) that was previously hypothesized to be responsible for allowing the ligand to enter the cavity and bind^44^ (**figure S8A, E**).

**Figure 5.**
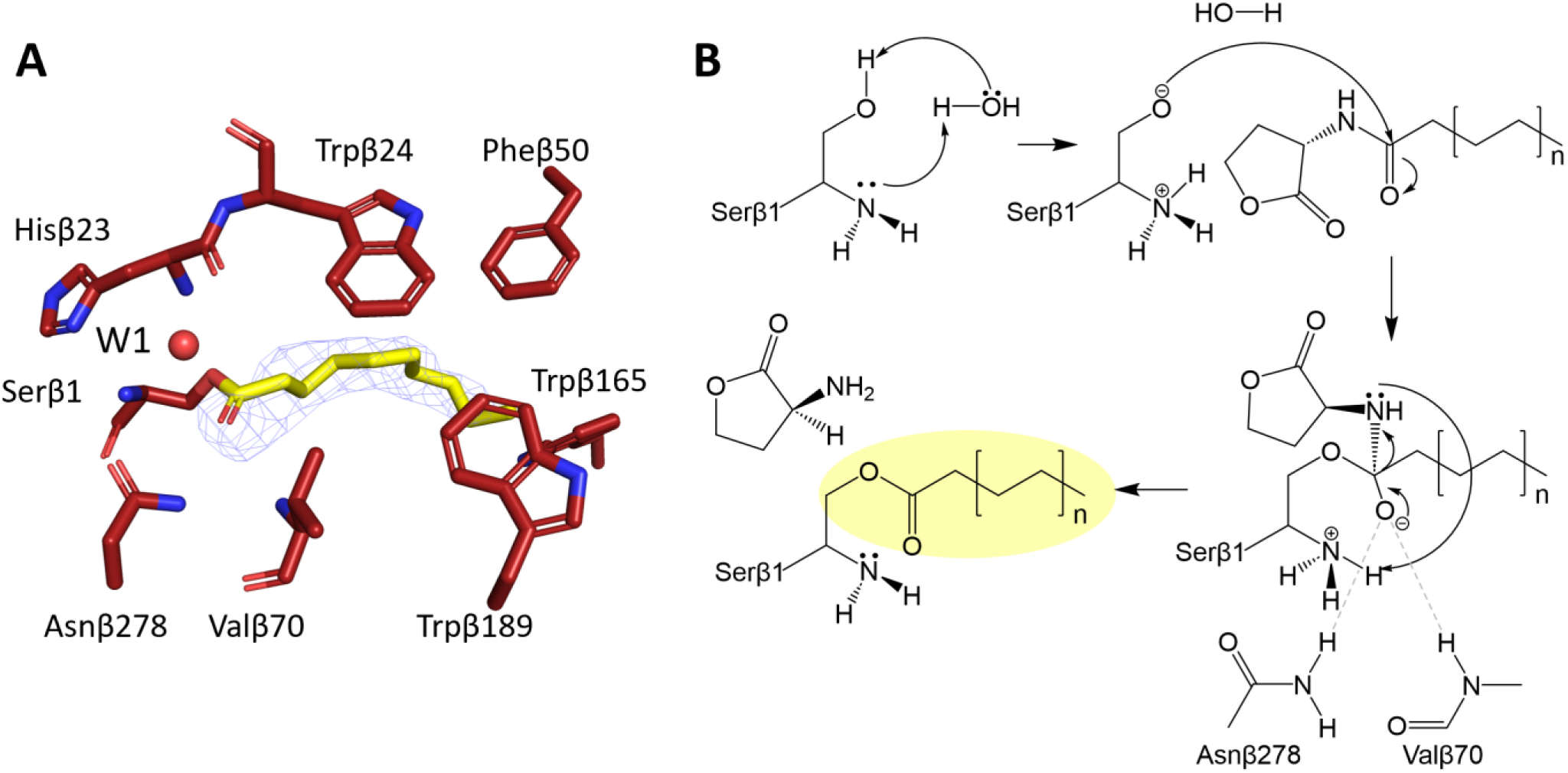
Acyl-enzyme intermediate shed lights on catalytic mechanism. **A**: Active site of M1. Blue mesh shows the Fourier difference Fo-Fc omit map contoured at 3.0 σ. W1: water. **B**: Reaction mechanism of M1 AHL-acylase with water as a base.

A previous report suggested that WT-MacQ may not utilize Serβ1 as the catalytic residue based on mutational data and biosensor-based activity measurements^54^. The capture of the acyl-enzyme intermediate in WT-MacQ^54^ and in the M1 structure supports the hypothesis that MacQ functions in the typical manner of Ntn-hydrolases^61,79^. In fact, the captured structure of M1 is compatible with the previously proposed mechanism for PvdQ^44^, where the free alpha-amino group of the catalytic Serβ1 acts as a general base to facilitate the activation of the serine nucleophile that attacks the carbonyl group of the AHL substrate. This generates a covalently bound, negatively charged, tetrahedral transition state that is stabilized by an oxyanion hole made of the Asnβ278 side chain and Valβ70 main chain NH groups. The developing negative charge of the transition states folds back and cause the departure of homoserine lactone. We note that the alpha-amino group of Serβ1 is in close vicinity (3-4 Å) to the leaving group and may give a proton (**figure 5B**), a role previously suggested to be performed by a second water molecule^44^. After the homoserine lactone release, the enzyme is acylated, a state illustrated by the obtained structure of M1 (**figure 5A**), and is likely recycled by a water molecule^44^.

Quorum quenching enzymes show considerable promise to contributing to the control of microbial behavior. Indeed, they were previously shown to reduce the virulence and biofilm formation of numerous bacteria. In order to use these enzymes as potential treatments or as an additive to improve materials, it appears important to source stable enzymes. Additionally, the complexity of quorum sensing may require enzymes that can operate selectively to target specific microbes or QS circuits, particularly in the context of communities where numerous signals are expected to be present and important.

In this context, the engineering of quorum quenching enzymes may contribute to the creation of more stable enzymes with altered kinetics. However, the engineering of AHL acylases is challenging because of the absence of convenient enzymatic assays. In fact, few studies on acylase engineering were produced, and reported improvements were significant but relatively modest. Here, we adapted an existing endpoint assay using OPA into a time-course assay, allowing for easy determination of AHL acylase kinetic parameters. We used the PROSS algorithm to generate variants for two acylase representatives, namely MacQ and PvdQ, and fully characterized these variants. We could obtain variants of PvdQ with significant increases in melting point temperature (up to 13.2 °C), and with superior ability to resist chemical solvents and formulation in coating. We also report variants of MacQ that exhibit >10-fold increase in catalytic efficiency against the short chain AHL substrates C4- and C6-HSL. Overall, the created variants show a unique combination of stability and activity levels. These changes in catalytic efficiency are translated into increased abilities to reduce virulence factors in the *P. aeruginosa* PA14 strain, with inhibitory levels increased by 78% for pyocyanin, 56% for elastase production, and 84% for protease activity as compared to the wild-type enzyme. In this effort, we were able to solve the crystal structure for one of the MacQ variants, M1. While this structure does not reveal obvious structural determinants explaining the observed changes in kinetics, it allowed for the capture of an acyl-enzyme intermediate that confirms the catalytic role of Serβ1 and suggest a similar catalytic mechanism to the previously proposed mechanism for WT-PvdQ^44^.

## METHODS

### Plasmid and sequence generation

Mutants for the reported *N-*acyl *L-*homoserine lactone acylase enzymes PvdQ^44,80^ and MacQ^53,54^ were generated using the Protein Repair One-Stop Shop (PROSS) algorithm^69^, which uses alignment scanning and computational mutation scanning^69^ to generate higher expressing and thermostable proteins. The chain B of the acylases, that contains the active site, was submitted to the server and PROSS outputs ten results with increasing proportion of mutations. In this study, we chose the top three results with the fewest mutations and characterized and compared them alongside their wild type counterparts. The sequences of the beta subunits were used as the input. The PDB structure 2WYE^44^ was used as the reference structure for PvdQ and 4YF9 was used for MacQ. The output generated can be found in supplemental files. The first three PROSS outputs for each protein were selected. These designed beta subunit sequences were combined with their original alpha subunit sequences minus the signal peptide sequences, as predicted by SignalP^81^. A TEV protease cleavage sequence (ENLYFQG) was inserted at the N-and C-termini of PvdQ and its mutants. Each sequence was ordered as a codon-optimized DNA insert between the *Nde*I and *Xho*I restriction sites in the pET-28a(+) vector from Twist Bioscience (San Francisco, CA, USA).

### Protein expression and purification

*Escherichia coli* Origami 2 (DE3) (Millipore Sigma, St. Louis, MO, USA) were transformed with pET-28a(+) vectors with inserts containing the proteins of interest to generate expression strains. Cells were grown in LB media supplemented with 50 μg/mL kanamycin and 10 μg/mL tetracycline at 37 °C. Protein expression was induced by the addition of 0.1-0.5 mM isopropyl ß-D-1-thiogalactopyranoside (IPTG), and cells were further incubated at 18 °C overnight. Cells were harvested by centrifugation at 6000 rpm for 10 minutes at 4 °C and the cell pellets stored at -20 °C until use. Lysis was performed by suspending the cell pellet in *acylase buffer* (50 mM HEPES pH 8.0, 150 mM sodium chloride, 10% glycerol) containing 1 mg/mL lysozyme, 2 mM phenylmethylsulfonyl fluoride (PMSF), and 1 μg/mL DNase, incubating on ice for 30 minutes, and subsequently sonicated for 1 minute using 1 s on and 2 s off intervals with a QSonica Q700 sonicator set to 45% amplitude. Ni-NTA resin (G-Biosciences, St. Louis, MO, USA) was added to the supernatant after clarification by centrifugation at 15 k rpm at 4 °C for 30 minutes, and binding was allowed to occur on a rotator at 4 °C for 1-2 hours. The lysate with resin was added to gravity columns. The resin was washed with acylase buffer containing 20 mM imidazole and the protein was eluted using acylase buffer containing 200 mM imidazole. Imidazole was removed from protein solutions through dialysis with acylase buffer, and protein was concentrated on centrifugal filters. Size exclusion chromatography was performed on a HiLoad 16/600, Superdex 200 pg column (GE Healthcare, Chicago, IL, USA) using acylase buffer without glycerol.

### Activity assays and biochemical characterization

Deacylation of *N-*acyl *L*-homoserine lactones was assayed by detecting the production of homoserine lactone using o-phthalaldehyde (OPA)^73^, similarly to previously reported fluorescence-based assays using fluorescamine^63,74^. Reaction mixture (200 μL) contained 1.9 mM OPA, 1 mM dithiothreitol (DTT), 5% dimethyl sulfoxide (DMSO) and 64-96 nM enzyme in 50 mM sodium borate pH 9.5. *N-*acyl *L-*homoserine lactone substrates (Cayman Chemical Company, Ann Arbor, MI, USA) were assayed at varying concentrations between 1 μM and 1 mM. The increase in fluorescence was measured at 37 °C in clear bottom 96-well black plates (Grenier Bio-one, Kremsmünster, Austria) at 360/460 nm. Kinetic parameters were determined by fitting the data to the Michaelis-Menten equation in GraphPad Prism 8, which also calculated the standard error values shown. When V_max_ could not be reached due to substrate solubility constraints or high K_M_ values, the catalytic efficiency was determined by fitting the data to a linear regression as appropriate. Samples with no enzyme or no substrate were used as the negative control. For solvent resistance, ethanol and ethyl acetate were added to the reaction mixture at the tested final percentage concentrations. Measurements were performed in quadruplicates. Outliers were excluded when reactions obviously failed or when experimental error was systematically introduced.

### Melting temperature assessment

Melting temperature of PvdQ and MacQ wild-type (WT) and mutants were determined using SYPRO Orange as the fluorogenic indicator of protein unfolding. 50 μL samples contained 15- 20 μg enzyme, 5x SYPRO Orange, and 50 mM sodium phosphate pH 7.4, 150 mM sodium chloride. The assay was carried out in triplicate in a 96-well PCR plate and measured in a qPCR thermocycler (Applied Biosystems, Waltham, MA). For enzymatic thermostability assays, enzyme solution was heated at 37, 40, 45, 50, 55 60, 65, 70, 75, and 80 °C in a thermocycler for 10 minutes. Samples were placed on ice for 5 minutes after heating then centrifuged at 15000 RCF for 5 minutes. The supernatant was then used in an endpoint OPA assay. Reactions contained 1.9 mM OPA, 1 mM DTT, 1.5 μg enzyme, and 0.2 mM C8-HSL in 50 mM sodium borate pH 9.5. Samples were incubated at 37 °C and the fluorescence at 360/460 nm measured after 30 minutes. The experiment was performed in triplicates. While both PvdQ and MacQ are multi-subunit enzymes, we did not observe a stepwise denaturation, nor did we see gradual loss of activity, indicating that the first disassociation step might render the entire oligomer inactive.

### Silicon paint tolerance assay

Silicon was mixed up at 10:1 ratio of Bluesil ESA 7246 A to Bluesil ESA 7246 B (Elsem Silicones, Oslo, Norway). Enzymes at 1 mg/mL in acylase buffer was mixed with silicon at a 1:10 ratio. Paint with buffer containing no enzyme was used as the background control. 10 μL of the enzyme-silicon mixture was spread on the bottom of wells in a clear flat bottom 96-well plate and placed on an oscillating shaker (VWR International, Radnor, PA, USA) for 5 minutes. Plates were allowed to dry and stored in the dark in a drawer before use. Acylase activity in the paint was assayed immediately or after 1, 7, and 21 days using the OPA assay containing 1.9 mM OPA, 1 mM DTT, and 0.1 mM 3-oxo-hexanoyl-L-homoserine lactone in 50 mM sodium borate pH 9.5 buffer at a total volume of 200 μL. The experiment was performed with five replicates.

### Virulence factor production assays

*Pseudomonas aeruginosa* strain PA14 (PA14) was obtained from Dr. Dianne Newmann at the California Institute of Technology and stored at -80 °C in 20% glycerol. PA14 was streaked on LB agar plates and incubated overnight at 37 °C. A single colony was used to inoculate 2 mL of LB and incubated with shaking for 4 hours or to an OD_600_ of 0.1. The preculture was diluted 1:100 with LB and 930 μL was added to sterile flat bottom 12-well plates to a final volume of 1 mL. 40 μL of enzymes diluted in 50 mM HEPES pH 7.8, 150 mM NaCl, 0.2 mM CoCl_2_ (PTE buffer) were added to each well at a final concentration of 50 μg/mL. Plates were sealed with BreatheEasy membrane (Diversified Biotech, Inc., Dedham, MA, USA), placed on a shaker, and incubated for 20 hours at 37 °C at 300 rpm. Cell growth was measured at 600 nm by diluting cultures by a factor of 20 in LB media. 1 mL of culture in each well was transferred into a 1.7 mL Eppendorf tube and pelleted at 14000 RCF for 10 minutes at room temperature. The supernatant was used for virulence factor production assays. Pyocyanin production was measured at 691 nm as previously described^20^ using 200 μL of culture supernatant in a clear, flat bottom 96-well plate. Elastase activity was measured using 50 μL of the cell supernatant in 250 μL reactions containing 5 mg/mL elastin-Congo red in 50 mM Tris-HCl pH 7.0, as previously described^20^. The reactions were incubated at 37 °C for 24 hours, then spun down at 2442 x rcf for 10 minutes. The supernatants were diluted by a factor of 10 in a clean 96-well plate and their absorbance measured at 491 nm. Protease activity was measured^20^ using 20 μL of the cell supernatant in 200 μL reactions containing 10 mg/mL azocasein in PBS pH 7.0. The reactions were incubated at 37 °C for 1 hour and the remaining substrate precipitated by the addition of 33.3 μL 20% trichloroacetic acid. The reactions were spun down at 2442 RCF for 10 minutes, the supernatants were transferred into a 96-well plate, and the absorbance measured at 366 nm. Absorbance values for virulence factor production assays were normalized against cell growth (OD 600 nm). Outliers were excluded when cells did not grow, overgrew, or were contaminated.

### Biosensor assays

A starter culture of *E. coli* JM109 containing pJBA132 carrying green fluorescent protein (GFP) under the control of LuxI/R was grown from a frozen glycerol stock overnight in 5 mL LB with 12.5 μg/mL tetracycline. The plasmid was obtained as a gift from Dr. David Daude (Gene&GreenTK, France). The starter culture was diluted 1:100 in LB media and incubated at 37 °C for 3 hours with shaking at 350 RPM. 50 μM of *N-*hexanoyl-*L-*homoserine lactone (C6- HSL) was incubated with 5 μg of acylase enzyme in 50 μL reactions of *acylase buffer* for 0, 20, 40, 60, 90, and 120 minutes at 37 °C. Reactions were stopped at those different timepoints by the addition of 50 μL DMSO, a concentration shown to inactivate the tested enzymes (data not shown). 2.5 μL of the terminated reaction was added to 197.5 μL of the biosensor culture in black clear bottom 96-well plates to induce expression of GFP and incubated further at 37 °C for 3 hours. DMSO was used as a background control and BSA was used as a negative control. The fluorescence of each well was measured at 485/20, 528/20 nm. Cell density was measured by absorbance at 600 nm and used to normalize GFP expression. The experiment was performed in quadruplicates.

### Crystallization

MacQ and its mutants were concentrated to 8 mg/mL in ultrafiltration units after size exclusion chromatography and crystallized in 100 mM Tris-HCl pH 6.5-8.5, 100 mM calcium acetate, and 14-18% polyethylene glycol (PEG) 3350. The best crystals were obtained in the lower pH range. Crystallization was allowed to occur at 18 °C through the hanging drop vapor diffusion method with drops at 1:1 and 1:2 precipitant:protein. Crystals began to appear 2+ months after drops were set up. Only crystals of variant M1 produced sufficiently large crystals.

### Data collection and structure determination

Crystals were transferred in a solution made of the mother liquor supplemented with 20% glycerol and flash cooled in liquid nitrogen. X-ray diffraction data was collected on the 23-ID-D beamline at the Advanced Photon Source (APS) in Argonne, Illinois (USA) using a wavelength of 1.0332 Å. Diffraction data and indexed, integrated, and scaled using the XDS software package^82^. M1 crystals were processed in the P1 space group (see **table S2**). Molecular replacement was performed using MOLREP^83^ using the MacQ structure (PDB: 4yf9^54^) as a model. Cycles of structure refinement and manual structure building was performed using REFMAC^84^ and Coot^85^. The M1 mutant structure was deposited into the Protein Data Bank under the accession code 8SO5.

## ACKNOWLEDGEMENTS

We are very grateful to the scientists at the Advanced Photon Source (APS, Argonne, IL, USA) and particularly the beamline scientists and coordinators at 23ID-D for their assistance.

## References

1. Fuqua, W. C., Winans, S. C. & Greenberg, E. P. Quorum sensing in bacteria: The LuxR-LuxI family of cell density-responsive transcriptional regulators. J. Bacteriol. 176, 269–275 (1994).

2. Case, R. J., Labbate, M. & Kjelleberg, S. AHL-driven quorum-sensing circuits: their frequency and function among the Proteobacteria. ISME J. 2, 345–349 (2008).

3. Smith, R. S. & Iglewski, B. H. Pseudomonas aeruginosa quorum sensing as a potential antimicrobial target. J. Clin. Invest. 112, 1460–1465 (2003).

4. Bhargava, N., Sharma, P. & Capalash, N. Quorum sensing in Acinetobacter: an emerging pathogen. Crit. Rev. Microbiol. 36, 349–360 (2010).

5. Eberl, L. Quorum sensing in the genus Burkholderia. Int. J. Med. Microbiol. 296, 103–110 (2006).

6. Suppiger, A., Schmid, N., Aguilar, C., Pessi, G. & Eberl, L. Two quorum sensing systems control biofilm formation and virulence in members of the *Burkholderia cepacia* complex. Virulence 4, 400–409 (2013).

7. McClean, K. H. et al. Quorum sensing and Chromobacterium violaceum:exploitation of violacein production and inhibition for the detection of N-acylhomoserine lactones.

8. Zhang, H.-B., Wang, L.-H. & Zhang, L.-H. Genetic control of quorum-sensing signal turnover in *Agrobacterium tumefaciens*. Proc. Natl. Acad. Sci. 99, 4638–4643 (2002).

9. Yeon, K.-M. et al. Quorum Sensing: A New Biofouling Control Paradigm in a Membrane Bioreactor for Advanced Wastewater Treatment. Environ. Sci. Technol. 43, 380– 385 (2009).

10. van Kessel, J. C., Ulrich, L. E., Zhulin, I. B. & Bassler, B. L. Analysis of Activator and Repressor Functions Reveals the Requirements for Transcriptional Control by LuxR, the Master Regulator of Quorum Sensing in Vibrio harveyi. mBio 4, e00378–13 (2013).

11. Winson, M. K. et al. Multiple N-acyl-L-homoserine lactone signal molecules regulate production of virulence determinants and secondary metabolites in Pseudomonas aeruginosa. Proc. Natl. Acad. Sci. 92, 9427–9431 (1995).

12. Bjarnsholt, T. et al. Pseudomonas aeruginosa tolerance to tobramycin, hydrogen peroxide and polymorphonuclear leukocytes is quorum-sensing dependent. Microbiology 151, 373–383 (2005).

13. Uroz, S. & Heinonsalo, J. Degradation of N-acyl homoserine lactone quorum sensing signal molecules by forest root-associated fungi. FEMS Microbiol. Ecol. 65, 271–278 (2008).

14. Khersonsky, O. & Tawfik, D. S. Structure−Reactivity Studies of Serum Paraoxonase PON1 Suggest that Its Native Activity Is Lactonase. Biochemistry 44, 6371–6382 (2005).

15. Draganov, D. I. et al. Human paraoxonases (PON1, PON2, and PON3) are lactonases with overlapping and distinct substrate specificities. J. Lipid Res. 46, 1239–1247 (2005).

16. Grandclément, C., Tannières, M., Moréra, S., Dessaux, Y. & Faure, D. Quorum quenching: role in nature and applied developments. FEMS Microbiol. Rev. 40, 86–116 (2016).

17. Dong, Y.-H. et al. Quenching quorum-sensing-dependent bacterial infection by an N-acyl homoserine lactonase. Nature 411, 813–817 (2001).

18. Migiyama, Y. et al. Efficacy of AiiM, an N-acylhomoserine lactonase, against pseudomonas aeruginosa in a mouse model of acute pneumonia. Antimicrob. Agents Chemother. 57, 3653–3658 (2013).

19. Hraiech, S. et al. Inhaled lactonase reduces pseudomonas aeruginosa quorum sensing and mortality in rat pneumonia. PLoS ONE 9, 1–8 (2014).

20. Mahan, K. et al. Effects of Signal Disruption Depends on the Substrate Preference of the Lactonase. Front. Microbiol. 10, 1–11 (2020).

21. Guendouze, A. et al. Effect of quorum quenching lactonase in clinical isolates of pseudomonas aeruginosa and comparison with quorum sensing inhibitors. Front. Microbiol. 8, 1–10 (2017).

22. Schwab, M. et al. Signal disruption leads to changes in bacterial community population. Front. Microbiol. 10, 1–13 (2019).

23. Mion, S. et al. Disrupting quorum sensing alters social interactions in Chromobacterium violaceum. NPJ Biofilms Microbiomes 7, 40 (2021).

24. Leadbetter, J. R. & Greenberg, E. P. Metabolism of acyl-homoserine lactone quorum-sensing signals by Variovorax paradoxus. J. Bacteriol. 182, 6921–6926 (2000).

25. Dong, Y.-H., Xu, J.-L., Li, X.-Z. & Zhang, L.-H. AiiA, an enzyme that inactivates the acylhomoserine lactone quorum-sensing signal and attenuates the virulence of Erwinia carotovora. (2000).

26. Tang, K. et al. MomL, a Novel Marine-Derived *N* -Acyl Homoserine Lactonase from Muricauda olearia. Appl. Environ. Microbiol. 81, 774–782 (2015).

27. Rémy, B. et al. Harnessing hyperthermostable lactonase from Sulfolobus solfataricus for biotechnological applications. Sci. Rep. 6, 1–11 (2016).

28. Huang, S., Bergonzi, C., Schwab, M., Elias, M. & Hicks, R. E. Evaluation of biological and enzymatic quorum quencher coating additives to reduce biocorrosion of steel. PLoS ONE 14, (2019).

29. Oh, H. S. et al. Control of membrane biofouling in MBR for wastewater treatment by quorum quenching bacteria encapsulated in microporous membrane. Environ. Sci. Technol. 46, 4877–4884 (2012).

30. Zhou, S., Zhang, A., Yin, H. & Chu, W. Bacillus sp. QSI-1 Modulate Quorum Sensing Signals Reduce Aeromonas hydrophila Level and Alter Gut Microbial Community Structure in Fish. Front. Cell. Infect. Microbiol. 6, (2016).

31. Kim, J.-H., Choi, D.-C., Yeon, K.-M., Kim, S.-R. & Lee, C.-H. Enzyme-Immobilized Nanofiltration Membrane To Mitigate Biofouling Based on Quorum Quenching. Environ. Sci. Technol. 45, 1601–1607 (2011).

32. de Celis, M. et al. Acylase enzymes disrupting quorum sensing alter the transcriptome and phenotype of Pseudomonas aeruginosa, and the composition of bacterial biofilms from wastewater treatment plants. Sci. Total Environ. 799, 149401 (2021).

33. Papaioannou, E. et al. Quorum-Quenching Acylase Reduces the Virulence of Pseudomonas aeruginosa in a Caenorhabditis elegans Infection Model. Antimicrob. Agents Chemother. 53, 4891–4897 (2009).

34. Koch, G. et al. Reducing virulence of the human pathogen Burkholderia by altering the substrate specificity of the quorum-quenching acylase PvdQ. Proc. Natl. Acad. Sci. 111, 1568–1573 (2014).

35. Ivanova, K. et al. Quorum-Quenching and Matrix-Degrading Enzymes in Multilayer Coatings Synergistically Prevent Bacterial Biofilm Formation on Urinary Catheters. ACS Appl. Mater. Interfaces 7, 27066–27077 (2015).

36. Grover, N. et al. Acylase-containing polyurethane coatings with anti-biofilm activity. Biotechnol. Bioeng. 113, 2535–2543 (2016).

37. Lee, J. et al. Immobilization and Stabilization of Acylase on Carboxylated Polyaniline Nanofibers for Highly Effective Antifouling Application via Quorum Quenching. ACS Appl. Mater. Interfaces 9, 15424–15432 (2017).

38. Utari, P. D., Setroikromo, R., Melgert, B. N. & Quax, W. J. PvdQ Quorum Quenching Acylase Attenuates Pseudomonas aeruginosa Virulence in a Mouse Model of Pulmonary Infection. Front. Cell. Infect. Microbiol. 8, (2018).

39. Ivanova, A., Ivanova, K., Tied, A., Heinze, T. & Tzanov, T. Layer-By-Layer Coating of Aminocellulose and Quorum Quenching Acylase on Silver Nanoparticles Synergistically Eradicate Bacteria and Their Biofilms. Adv. Funct. Mater. 30, (2020).

40. Vogel, J., Wakker-Havinga, M., Setroikromo, R. & Quax, W. J. Immobilized Acylase PvdQ Reduces Pseudomonas aeruginosa Biofilm Formation on PDMS Silicone. Front. Chem. 8, 1–9 (2020).

41. Park, S.-Y., et al. *N* -acylhomoserine lactonase producing *Rhodococcus* spp. with different AHL-degrading activities. FEMS Microbiol. Lett. 261, 102–108 (2006).

42. Koch, G., Nadal-Jimenez, P., Cool, R. H. & Quax, W. J. Deinococcus radiodurans can interfere with quorum sensing by producing an AHL-acylase and an AHL-lactonase. FEMS Microbiol. Lett. 356, 62–70 (2014).

43. Yates, E. A. et al. N-acylhomoserine lactones undergo lactonolysis in a pH-, temperature-, and acyl chain length-dependent manner during growth of Yersinia pseudotuberculosis and Pseudomonas aeruginosa. Infect. Immun. 70, 5635–5646 (2002).

44. Bokhove, M., Jimenez, P. N., Quax, W. J. & Dijkstra, B. W. The quorum-quenching N-acyl homoserine lactone acylase PvdQ is an Ntn-hydrolase with an unusual substrate-binding pocket. Proc. Natl. Acad. Sci. 107, 686–691 (2010).

45. Mukherji, R., Varshney, N. K., Panigrahi, P., Suresh, C. G. & Prabhune, A. A new role for penicillin acylases: Degradation of acyl homoserine lactone quorum sensing signals by Kluyvera citrophila penicillin G acylase. Enzyme Microb. Technol. 56, 1–7 (2014).

46. Sunder, A. V. et al. Penicillin V acylases from gram-negative bacteria degrade N-acylhomoserine lactones and attenuate virulence in Pseudomonas aeruginosa. Appl. Microbiol. Biotechnol. 101, 2383–2395 (2017).

47. Utari, P. D., Vogel, J. & Quax, W. J. Deciphering Physiological Functions of AHL Quorum Quenching Acylases. Front. Microbiol. 8, 1123 (2017).

48. Sio, C. F. et al. Quorum quenching by an N-acyl-homoserine lactone acylase from Pseudomonas aeruginosa PAO1. Infect. Immun. 74, 1673–1682 (2006).

49. Huang, J. J., Petersen, A., Whiteley, M. & Leadbetter, J. R. Identification of QuiP, the Product of Gene PA1032, as the Second Acyl-Homoserine Lactone Acylase of Pseudomonas aeruginosa PAO1. Appl. Environ. Microbiol. 72, 1190–1197 (2006).

50. Jimenez, P. N. et al. Role of PvdQ in Pseudomonas aeruginosa virulence under iron-limiting conditions. Microbiology 156, 49–59 (2010).

51. Clevenger, K. D., Wu, R., Er, J. A. V., Liu, D. & Fast, W. Rational Design of a Transition State Analogue with Picomolar Affinity for Pseudomonas aeruginosa PvdQ, a Siderophore Biosynthetic Enzyme. ACS Chem. Biol. 8, 2192–2200 (2013).

52. Maisuria, V. B. & Nerurkar, A. S. Interference of quorum sensing by Delftia sp. VM4 depends on the activity of a novel n-acylhomoserine lactone-acylase. PLoS ONE 10, 1–15 (2015).

53. Kusada, H., Tamaki, H., Kamagata, Y., Hanada, S. & Kimura, N. A novel quorumquenching N-acylhomoserine lactone acylase from Acidovorax sp. strain MR-S7 mediates antibiotic resistance. Appl. Environ. Microbiol. 83, 1–9 (2017).

54. Yasutake, Y. et al. Bifunctional quorum-quenching and antibiotic-acylase MacQ forms a 170-kDa capsule-shaped molecule containing spacer polypeptides. Sci. Rep. 7, 1–11 (2017).

55. Wellington, S. & Greenberg, E. P. Quorum Sensing Signal Selectivity and the Potential for Interspecies Cross Talk. mBio 10, 1–22 (2019).

56. Kim, J.-H., Lee, S.-C., Kyeong, H.-H. & Kim, H.-S. A Genetic Circuit System Based on Quorum Sensing Signaling for Directed Evolution of Quorum-Quenching Enzymes. ChemBioChem 11, 1748–1753 (2010).

57. Hiblot, J., Gotthard, G., Elias, M. & Chabriere, E. Differential Active Site Loop Conformations Mediate Promiscuous Activities in the Lactonase SsoPox. PLoS ONE 8, 1–14 (2013).

58. Billot, R. et al. Engineering acyl-homoserine lactone-interfering enzymes toward bacterial control. J. Biol. Chem. 295, 12993–13007 (2020).

59. Billot, R. et al. Applying molecular and phenotypic screening assays to identify efficient quorum quenching lactonases. Enzyme Microb. Technol. 160, 110092 (2022).

60. Charlton, T. S. et al. A novel and sensitive method for the quantification of N-3- oxoacyl homoserine lactones using gas chromatography-mass spectrometry: Application to a model bacterial biofilm. Environ. Microbiol. 2, 530–541 (2000).

61. Lin, Y.-H. et al. Acyl-homoserine lactone acylase from Ralstonia strain XJ12B represents a novel and potent class of quorum-quenching enzymes. Mol. Microbiol. 47, 849– 860 (2003).

62. Huang, J. J., Han, J.-I., Zhang, L.-H. & Leadbetter, J. R. Utilization of Acyl-Homoserine Lactone Quorum Signals for Growth by a Soil Pseudomonad and Pseudomonas aeruginosa PAO1. Appl. Environ. Microbiol. 69, 5941–5949 (2003).

63. Murugayah, S. A., Warring, S. L. & Gerth, M. L. Optimisation of a high-throughput fluorescamine assay for detection of N-acyl-L-homoserine lactone acylase activity. Anal. Biochem. 566, 10–12 (2019).

64. Wahjudi, M. et al. PA0305 of pseudomonas aeruginosa is a quorum quenching acylhomoserine lactone acylase belonging to the Ntn hydrolase superfamily. Microbiology 157, 2042–2055 (2011).

65. Winson, M. K. et al. Construction and analysis of *luxCDABE* -based plasmid sensors for investigating *N* -acyl homoserine lactone-mediated quorum sensing. FEMS Microbiol. Lett. 163, 185–192 (1998).

66. Steindler, L. & Venturi, V. Detection of quorum-sensing N-acyl homoserine lactone signal molecules by bacterial biosensors. FEMS Microbiol. Lett. 266, 1–9 (2007).

67. Romero, M., Diggle, S. P., Heeb, S., Cámara, M. & Otero, A. Quorum quenching activity in Anabaena sp. PCC 7120: Identification of AiiC, a novel AHL-acylase. FEMS Microbiol. Lett. 280, 73–80 (2008).

68. Andersen, J. B., et al. *gfp* -Based *N* -Acyl Homoserine-Lactone Sensor Systems for Detection of Bacterial Communication. Appl. Environ. Microbiol. 67, 575–585 (2001).

69. Goldenzweig, A. et al. Automated Structure-and Sequence-Based Design of Proteins for High Bacterial Expression and Stability. Mol. Cell 63, 337–346 (2016).

70. Duda, C. T. & Kissinger, P. T. Determination of Biogenic Amines, Their Metabolites, and Other Neurochemicals by Liquid Chromatography/Electrochemistry. Handb. Behav. Neurosci. 11, 41–82 (1993).

71. Morohoshi, T., Nakazawa, S., Ebata, A., Kato, N. & Ikeda, T. Identification and characterization of N-acylhomoserine lactone-acylase from the fish intestinal Shewanella sp. strain MIB015. Biosci. Biotechnol. Biochem. 72, 1887–1893 (2008).

72. Islam, M. M., Kobayashi, K., Kidokoro, S. & Kuroda, Y. Hydrophobic surface residues can stabilize a protein through improved water–protein interactions. FEBS J. 286, 4122–4134 (2019).

73. Xu, F., Byun, T., Dussen, H.-J. & Duke, K. R. Degradation of N-acylhomoserine lactones, the bacterial quorum-sensing molecules, by acylase. J. Biotechnol. 101, 89–96 (2003).

74. Reyes, F., Martinez, M.-J & Soliveri, J. Determination of cephalosporin-C amidohydrolase activity with fluorescamine. J. Pharm. Pharmacol. 41, 136–137 (1989).

75. Roth, Marc. Fluorescence reaction for amino acids. Anal. Chem. 43, 880–882 (1971).

76. Lee, J. & Zhang, L. The hierarchy quorum sensing network in Pseudomonas aeruginosa. Protein Cell 6, 26–41 (2015).

77. Bergonzi, C., Schwab, M., Naik, T. & Elias, M. The Structural Determinants Accounting for the Broad Substrate Specificity of the Quorum Quenching Lactonase GcL. ChemBioChem 20, 1848–1855 (2019).

78. Rémy, B. et al. Lactonase Specificity Is Key to Quorum Quenching in Pseudomonas aeruginosa. Front. Microbiol. 11, 1–17 (2020).

79. Duggleby, H. J. et al. Penicillin acylase has a single-amino-acid catalytic centre. Nature 373, 264–268 (1995).

80. Sio, C. F. et al. Quorum quenching by an N-acyl-homoserine lactone acylase from Pseudomonas aeruginosa PAO1. Infect. Immun. 74, 1673–1682 (2006).

81. Almagro Armenteros, J. J., et al. SignalP 5.0 improves signal peptide predictions using deep neural networks. Nat. Biotechnol. 37, 420–423 (2019).

82. Kabsch, W. XDS. Acta Crystallogr. D Biol. Crystallogr. 66, 125–132 (2010).

83. Vagin, A. & Teplyakov, A. Molecular replacement with MOLREP. Acta Crystallogr. D Biol. Crystallogr. 66, 22–25 (2010).

84. Murshudov, G. N. et al. REFMAC5 for the refinement of macromolecular crystal structures. Acta Crystallogr. D Biol. Crystallogr. 67, 355–367 (2011).

85. Emsley, P., Lohkamp, B., Scott, W. G. & Cowtan, K. Features and development of Coot. Acta Crystallogr. D Biol. Crystallogr. 66, 486–501 (2010).

