## Supplementary Material for "Engineering Quorum Quenching Acylases with Improved Kinetic and Biochemical Properties"

**Supporting information for "Engineering Quorum Quenching Acylases with Improved Kinetic and Biochemical Properties"**

**Summary**


**Figure S1**: SDS-PAGE analysis of mutant acylase variants after purification. 3

**Figure S2**: Acylase activity against C8-HSL after heat treatment. 4

**Figure S3**: Schematic of real-time AHL acylase assay. 5

**Figure S4**: Real-time measurement of acylase activity. 6

**Figure S5**: Kinetics for WT-PvdQ and variants used in this study. 7

**Figure S6**: Kinetics for WT-MacQ and variants used in this study. 8

**Figure S7:** Examples of conformational changes observed between M1 and wild-type MacQ enzyme structure. 9

**Figure S8**: Comparison of different AHL acylase active sites. 10

**Table S1**: Amino acid substitutions in the acylase variants used in this study. 11

**Table S2**: Amino acid substitutions and their properties for MacQ variants. 13

**Table S3**: Kinetic parameters for wild-type PvdQ acylase from *Pseudomonas aeruginosa*. 14

**Table S4**: Data collection and refinement statistics of the M1 structure. 15

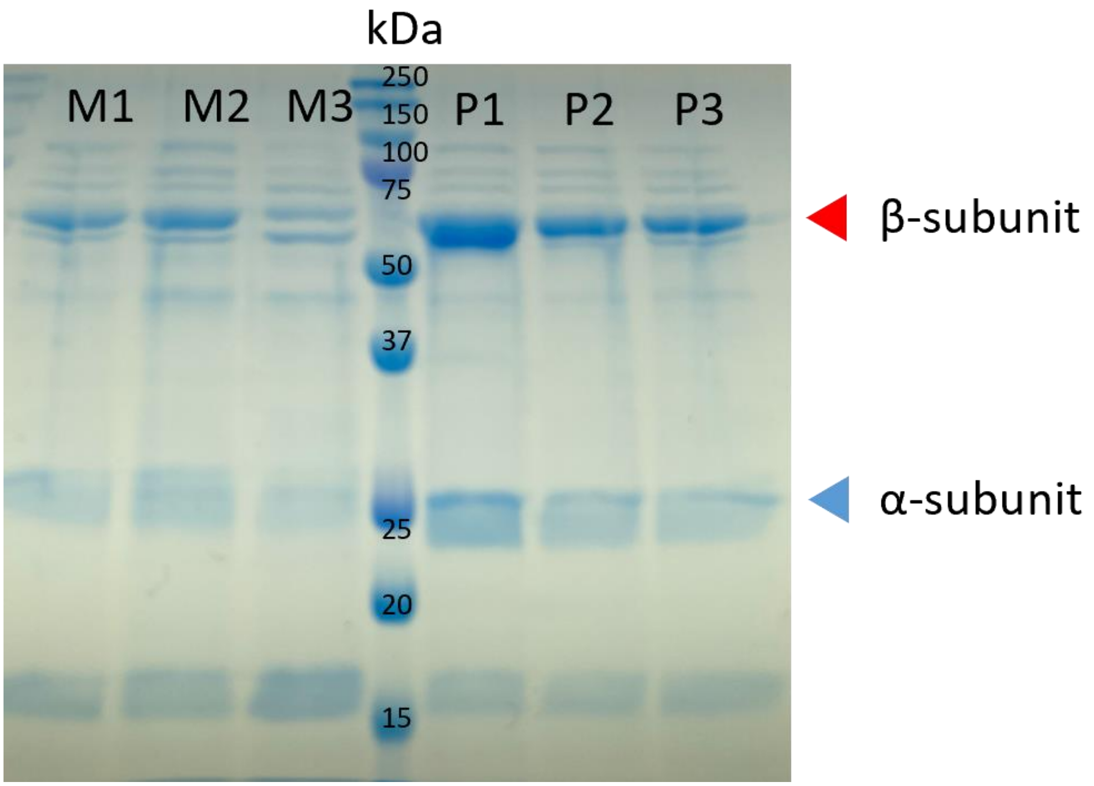


**Figure S1**: SDS-PAGE analysis of acylase variants after purification through IMAC. Higher molecular weight band may correspond to an uncleaved proenzyme.

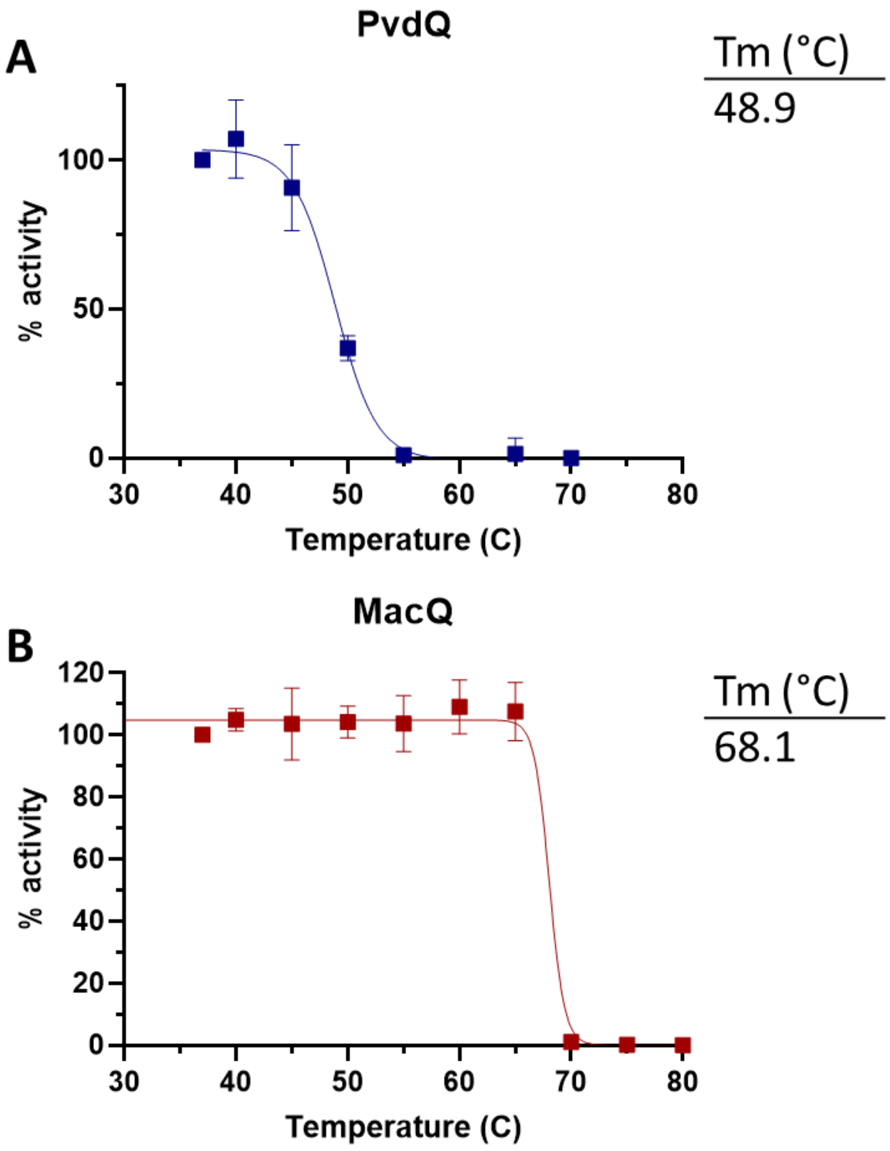


**Figure S2**: Acylase activity against C8-HSL after heat treatment as measured by OPA assay as a fraction of original activity. Error bars show standard deviation. N=3.

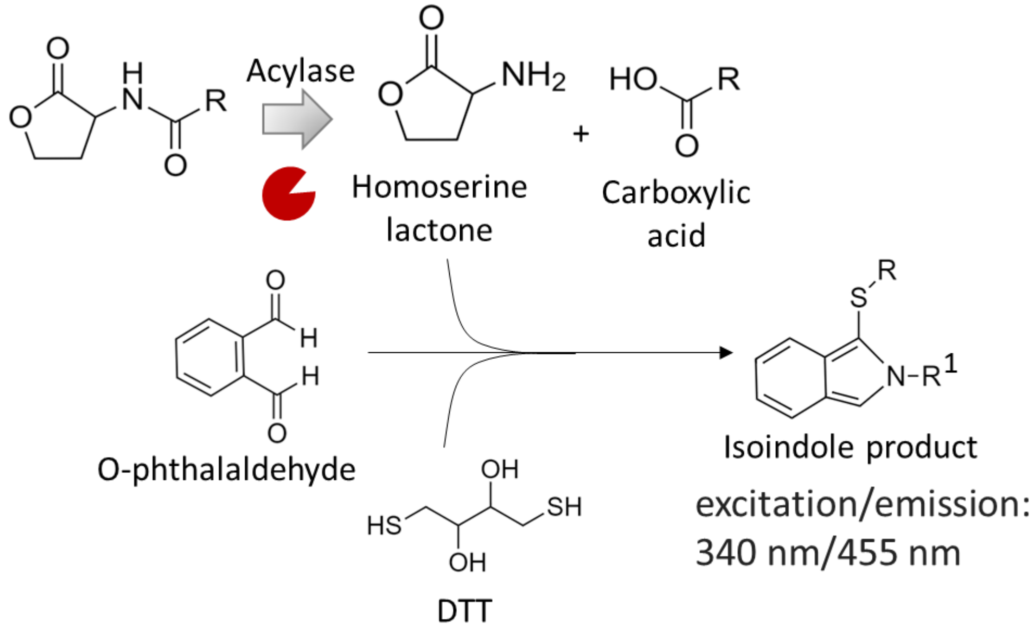


**Figure S3**: Schematic of reaction used to monitor the acylase activity. AHL acylases cause the release of homoserine lactone and carboxylic acid. The homoserine lactone reacts with o-phthalaldehyde (OPA) and dithiothreitol (DTT) to form a fluorescent isoindole product.

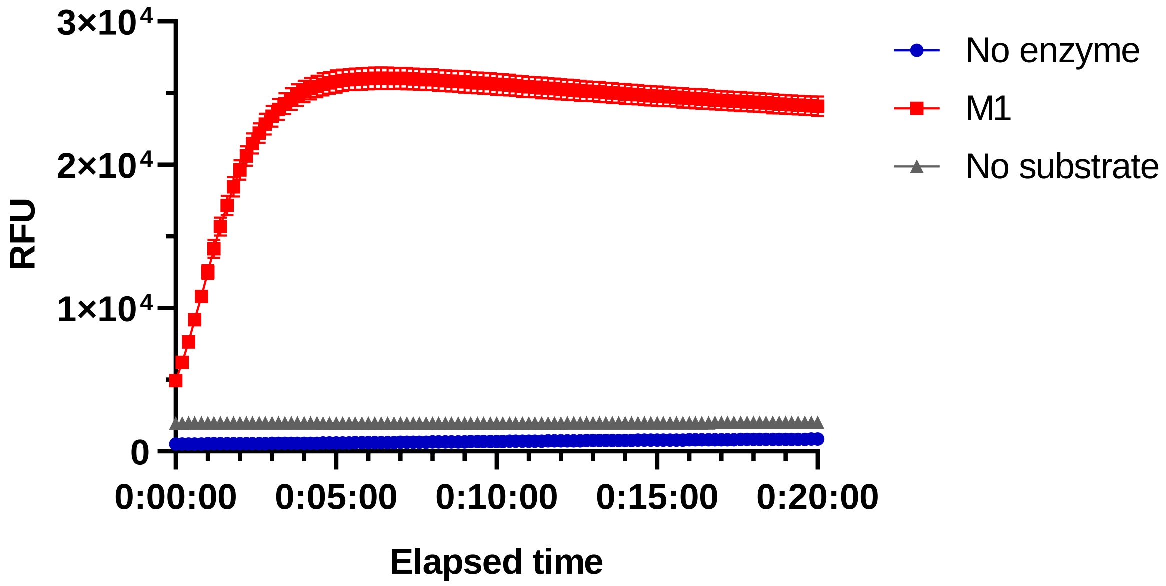


**Figure S4**: Real-time increase in fluorescence using the OPA assay to measure AHL acylase activity. The substrate used is *N*-decanoyl-L-homoserine lactone at 0.1 mM final concentration. 2 µg of M1 was used. DMSO was used as the negative control in the no substrate sample controls (grey) and enzyme buffer was used in the no enzyme sample controls (blue). Error bars show standard deviation. Measurements were repeated in triplicate. RFU: relative fluorescence units.

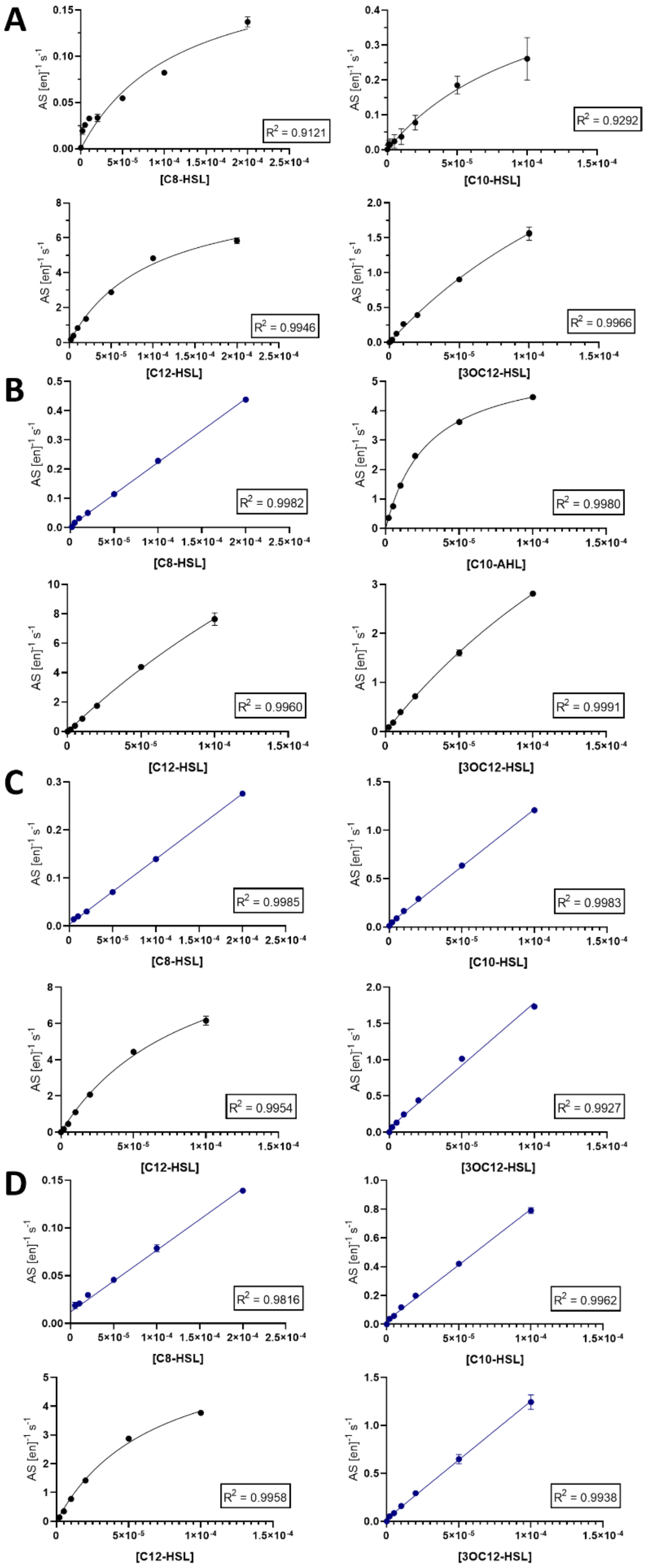


**Figure S5**: Initial rates of reaction of PvdQ and variants against varying concentrations of AHL substrates. Curves are fitted to Michaelis-Menten (black) or as linear regressions (blue) as appropriate. **A**: PvdQ **B**: P1 **C**: P2 **D**: P3. Substrate concentrations are in Mol. N=3-4. Error bars show standard deviation.

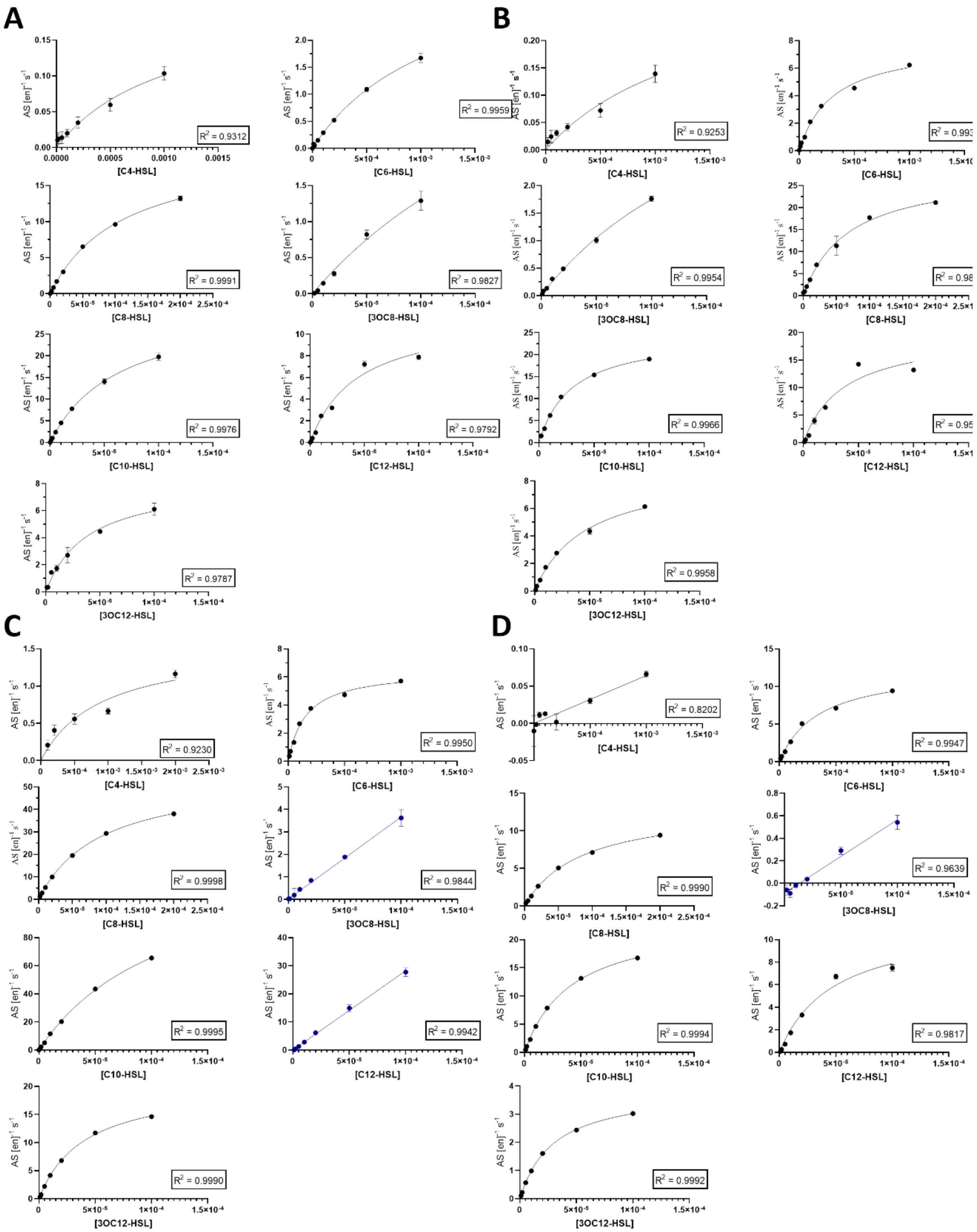


**Figure S6**: Initial rates of reactions of MacQ and variants against varying concentrations of AHL substrates. Curves are fitted to the Michaelis-Menten equation (black) or as linear regressions (blue) as appropriate. **A**: MacQ **B**: M1 **C**: M2 **D**: M3. Substrate concentrations are in Mol. N=3-4. Error bars show standard deviation.

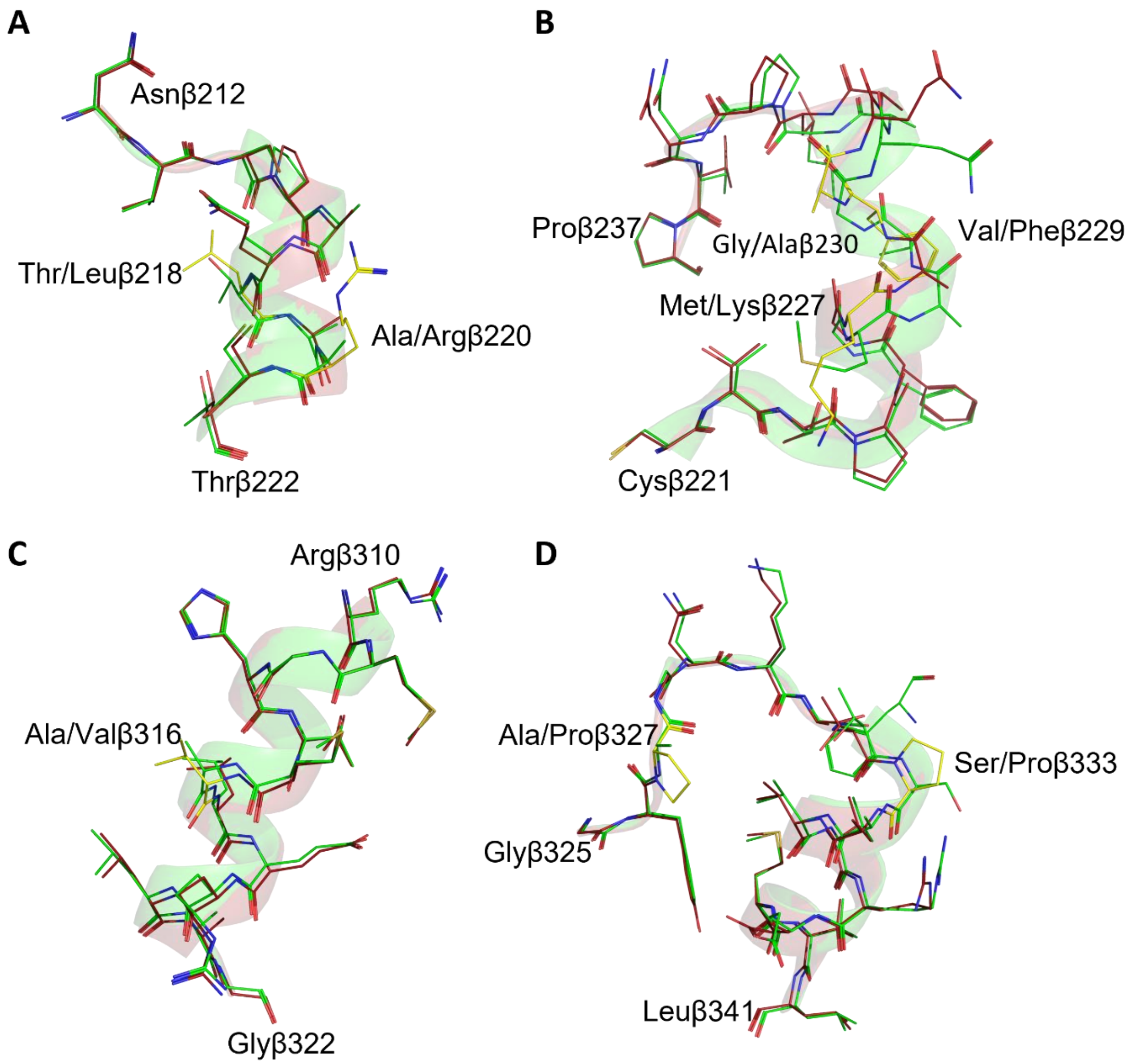


**Figure S7:** Examples of conformational changes observed between M1 and wild-type MacQ enzyme structure (PDB: 4yfa). Mutation sites are showed as yellow sticks. **A**: region between Asnβ212 and Thrβ222 containing Thrβ218Leu and Alaβ220Arg mutations. **B**: Region between Cysβ221 and Proβ237 containing Metβ227Lys, Valβ229Phe, and Glyβ230Ala mutations. **C**: Region between Argβ310 and Gluβ322 containing an Alaβ316Val mutation. **D**: Region between Glyβ325 and Leuβ341 containing Alaβ327Pro and Serβ333Pro mutations.

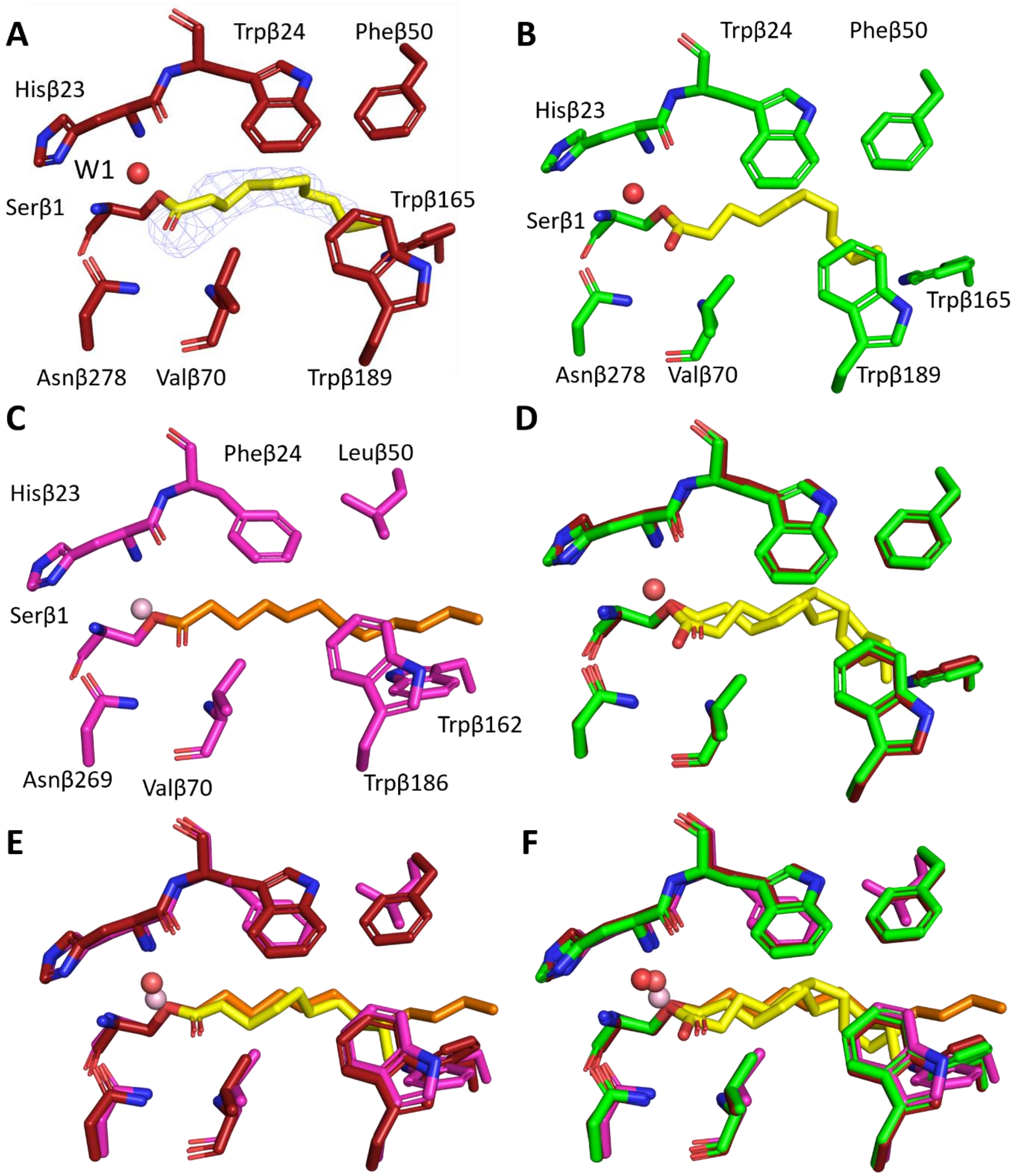


**Figure S8**: Comparison of the AHL acylase active sites. **A**: M1 (maroon, PDB: 8so5). **B**: WT-MacQ (green, PDB: 4yfa). **C**: WT-PvdQ (magenta, PDB: 2wyb). **D**: M1 (maroon) and WT-MacQ (green) superimposed. **E**: M1 (maroon) and WT-PvdQ (magenta) superimposed. **F**: M1 (maroon), WT-MacQ (green), and WT-PvdQ (magenta) are all superimposed. Decanoic acid ligand in yellow and dodecanoic acid ligand in orange.

**Table S1**: Amino acid substitutions in the acylase variants used in this study.

| Position | PvdQ | P1 | P2 | P3 |
|---|---|---|---|---|
| 61 | S | N | N | N |
| 82 | A | A | A | Q |
| 100 | L | M | M | M |
| 104 | S | S | T | T |
| 127 | I | R | R | R |
| 194 | E | R | R | R |
| 203 | Q | V | V | I |
| 209 | L | V | V | V |
| 210 | K | K | T | T |
| 214 | I | L | L | L |
| 215 | P | Q | Q | Q |
| 233 | Q | Q | T | T |
| 238 | A | D | D | D |
| 244 | A | R | R | R |
| 248 | A | A | P | P |
| 251 | T | V | V | V |
| 327 | S | S | S | A |
| 361 | G | R | T | T |
| 368 | S | M | M | M |
| 377 | M | W | W | W |
| 389 | E | V | V | V |
| 393 | A | P | P | P |
| 401 | Q | R | R | R |
| 433 | G | D | D | D |
| 446 | Q | D | D | D |
| 450 | A | P | P | P |
| 500 | F | F | F | Y |

| Position | MacQ | M1 | M2 | M3 |
|---|---|---|---|---|
| 17 | V | L | L | L |
| 63 | S | S | H | H |
| 82 | S | S | S | Q |
| 85 | Q | P | P | P |
| 87 | E | D | D | D |
| 93 | R | V | V | V |
| 146 | G | Q | Q | Q |
| 186 | S | G | A | A |
| 199 | S | D | D | D |
| 209 | A | V | V | V |
| 218 | T | L | L | L |
| 220 | A | R | R | R |
| 227 | M | K | K | K |
| 229 | V | F | F | F |
| 230 | G | A | A | A |
| 257 | K | P | P | P |
| 260 | V | I | I | I |
| 316 | A | V | V | V |
| 317 | L | L | Q | Q |
| 327 | A | P | P | P |
| 333 | S | P | P | P |
| 342 | G | G | G | D |
| 343 | S | N | N | N |
| 371 | A | A | E | E |
| 395 | S | A | A | A |
| 439 | M | M | L | L |
| 443 | I | V | V | V |

| | | | | |
|---|---|---|---|---|
| 529 | S | S | T | T |
| 534 | D | D | K | K |
| # mutations | 0 | 20 | 26 | 29 |
| % mutations | 0 | 3.66 | 4.76 | 5.31 |

| | | | | |
|---|---|---|---|---|
| 447 | G | G | R | R |
| 451 | Y | Y | Y | F |
| 463 | A | V | V | V |
| 527 | F | Y | Y | Y |
| 539 | G | G | T | T |
| # mutations | 0 | 23 | 29 | 32 |
| % mutations | 0 | 4 | 5.04 | 5.57 |

**Table S2**: Amino acid substitutions and their properties for MacQ variants.

| Position | MacQ | M1 | M2 | M3 |
|---|---|---|---|---|
| 260 | V | I | I | I |
| 229 | V | F | F | F |
| 17 | V | L | L | L |
| 218 | T | L | L | L |
| 395 | S | A | A | A |
| 343 | S | N | N | N |
| 333 | S | P | P | P |
| 199 | S | D | D | D |
| 186 | S | G | A | A |
| 93 | R | V | V | V |
| 85 | Q | P | P | P |
| 227 | M | K | K | K |
| 257 | K | P | P | P |
| 443 | I | V | V | V |
| 230 | G | A | A | A |
| 146 | G | Q | Q | Q |
| 527 | F | Y | Y | Y |
| 87 | E | D | D | D |
| 463 | A | V | V | V |
| 327 | A | P | P | P |
| 316 | A | V | V | V |
| 220 | A | R | R | R |
| 209 | A | V | V | V |
| 63 | S | | H | H |
| 439 | M | | L | L |
| 317 | L | | Q | Q |
| 539 | G | | T | T |
| 447 | G | | R | R |
| 371 | A | | E | E |
| 451 | Y | | | F |
| 82 | S | | | Q |
| 342 | G | | | D |

- polar uncharged
- hydrophobic
- positively charged
- negatively charged
- uncharged

**Table S3**: Kinetic parameters for wild-type PvdQ acylase from *Pseudomonas aeruginosa*.

| | This study | | | Clevenger et al., 2013[51] | | | Koch et al., 2014[34] | | |
|---|---|---|---|---|---|---|---|---|---|
| HSL | $K_{cat}$ ($s^{-1}$) | $K_M$ (μM) | $K_{cat}/K_M$ ($s^{-1}$ $M^{-1}$) | $K_{cat}$ ($s^{-1}$) | $K_M$ (μM) | $K_{cat}/K_M$ ($s^{-1}$ $M^{-1}$) | $K_{cat}$ ($s^{-1}$) | $K_M$ (μM) | $K_{cat}/K_M$ ($s^{-1}$ $M^{-1}$) |
| C4 | na | na | na | NR | NR | NR | NR | NR | NR |
| C6 | na | na | na | NR | NR | NR | NR | NR | NR |
| C8 | (2.06 ± 0.30) x $10^{-1}$ | (1.18 ± 0.34) x $10^2$ | (1.75 ± 0.57) x $10^3$ | ND | ND | 2.20 x $10^2$ | NR | NR | 0.80 x $10^3$ |
| C10 | (5.63 ± 1.29) x $10^{-1}$ | (1.13 ± 0.42) x $10^2$ | (5.00 ± 2.17) x $10^3$ | ND | ND | 2.20 x$10^3$ | NR | NR | NR |
| C12 | 8.91 ± 0.32 | 98.1 ± 34.1 | (9.08 ± 0.76) x $10^4$ | 2.47 ± 0.117 | 11.0 ± 2 | 2.20 x $10^5$ | NR | NR | NR |
| 3OC12 | 5.23 ± 0.55 | (2.35 ± 0.07) x $10^2$ | (2.22 ± 0.40) x $10^4$ | ND | ND | 2.30 x $10^3$ | NR | NR | 5.80 x $10^3$ |

na: no activity
NR: not reported
ND: not determined due to substrate solubility
C4: *N*-butyryl-L-homoserine lactone. C6: *N*-hexanoyl-L-homoserine lactone. C8: *N*-octanoyl-L-homoserine lactone. C10: *N*-decanoyl-L-homoserine lactone. C12: *N*-dodecanoyl-L-homoserine-lactone. 3OC12: 3-oxo-dodecanoyl-L-homoserine lactone.

**Table S4**: Data collection and refinement statistics of the M1 structure.

| Data collection statistics | MacQ-M1 |
|---|---|
| PDB ID | 8SO5 |
| Resolution (Å) | 2.35 |
| Diffraction source | APS Argonne 23ID-D |
| Wavelength (Å) | 1.0332 |
| Detector | Dectris Pilatus3-6M |
| Rotation range per image (°) | 0.5 |
| Total rotation range (°) | 165 |
| Space group | $P2_1$ |
| Unit-cell parameters (Å) | a = 97.07, b = 80.65, c = 101.60; α= 90.000 γ= 90.000, β= 90.814 |
| Resolution range (Å) | 2.35 (2.45-2.35) |
| N° of reflections (last bin) | 200867 (23308) |
| N° of unique reflections (last bin) | 64842 (7609) |
| Completeness (%) (last bin) | 98.9 (99.3) |
| Redundancy (last bin) | 3.10 (3.06) |
| $\langle I/\sigma(I)\rangle$ (last bin) | 11.76 (2.92) |
| $R_{meas}$ (%) (last bin) | 10.9 (56.1) |
| $CC_{1/2}$ (last bin) | 99.4 (81.1) |
| **Refinement statistics** | |
| N° of total model atoms | 11919 |
| $R_{free}/R_{work}$ | 22.88/18.60 |
| Ramachandran core and allowed (%) | 87.5 |
| Ramachandran generously allowed (%) | 12.0 |
| Ramachandran outliers (%) | 0.5 |
| *Rmsd from ideal* | |
| Bond lengths (Å) | 0.001 |
| Bond angles (°) | 0.582 |
